# Hierarchical complexity of the macro-scale neonatal brain

**DOI:** 10.1101/2020.01.16.909150

**Authors:** Manuel Blesa, Paola Galdi, Simon R. Cox, Gemma Sullivan, David Q. Stoye, Gillian J. Lamb, Alan J. Quigley, Michael J. Thrippleton, Javier Escudero, Mark E. Bastin, Keith M. Smith, James P. Boardman

**Affiliations:** MRC Centre for Reproductive Health, University of Edinburgh, Edinburgh EH16 4TJ, UK; Lothian Birth Cohorts group, Department of Psychology, University of Edinburgh, Edinburgh EH8 9JZ, UK; Department of Radiology, Royal Hospital for Sick Children, Edinburgh EH9 1LF, UK; Centre for Clinical Brain Sciences, University of Edinburgh, Edinburgh EH16 4SB, UK; Edinburgh Imaging, University of Edinburgh, Edinburgh EH16 4SB, UK; School of Engineering, Institute for Digital Communications, University of Edinburgh, Edinburgh EH9 3FG, UK; Usher Institute, University of Edinburgh, Edinburgh EH16 4UX, UK; Health Data Research UK, London NW1 2BE, UK

**Keywords:** Hierarchical Complexity, Newborn, Developing Brain, Network Analysis, Structural Connectome, dMRI

## Abstract

The human adult structural connectome has a rich nodal hierarchy, with highly diverse connectivity patterns aligned to the diverse range of functional specializations in the brain. The emergence of this hierarchical complexity in human development is unknown. Here, we substantiate the hierarchical tiers and hierarchical complexity of brain networks in the newborn period; assess correspondences with hierarchical complexity in adulthood; and investigate the effect of preterm birth, a leading cause of atypical brain development and later neurocognitive impairment, on hierarchical complexity. We report that neonatal and adult structural connectomes are both composed of distinct hierarchical tiers, and that hierarchical complexity is greater in term born neonates than in preterms. This is due to diversity of connectivity patterns of regions within the intermediate tiers, which consist of regions that underlie sensorimotor processing and its integration with cognitive information. For neonates and adults, the highest tier (hub regions) is ordered, rather than complex, with more homogeneous connectivity patterns in structural hubs. This suggests that the brain develops first a more rigid structure in hub regions allowing for the development of greater and more diverse functional specialization in lower level regions, while connectivity underpinning this diversity is dysmature in infants born preterm.

## Introduction

The integrity of brain development during pregnancy and the newborn period is critical for life long cognitive function and brain health. During the second and third trimesters of pregnancy, there is a phase of rapid brain maturation characterized by volumetric growth, increases in cortical complexity, white matter (WM) organization and myelination (Boardman and Counsell, 2019). Early exposure to extrauterine life due to preterm birth affects around 11% of births, and is closely associated with neurodevelopmental, cognitive and psychiatric impairment (Nosarti et al., 2012; Mathewson et al., 2017; Wolke et al., 2019), and alterations to development (Batalle et al., 2018b) that are apparent using *in vivo* imaging techniques. At the macro scale, these alterations can be characterized by charting WM connections between brain regions using diffusion MRI (dMRI) (Ball et al., 2013; Van Den Heuvel et al., 2015; Brown et al., 2014; Batalle et al., 2017; Zhao et al., 2019a; Lee et al., 2019; Galdi et al., 2020).

The resulting structural brain network– or connectome– has been extensively explored using popular network metrics in the neonatal brain. Findings indicate similar characteristics as found ubiquitously in real-world networks, including local and global efficiency, high clustering coefficient and short characteristic path length (i.e. small-worldness), and a strong rich club coefficient (Van Den Heuvel et al., 2015; Brown et al., 2014; Batalle et al., 2017; Lee et al., 2019; Ball et al., 2014). Often they reveal remarkable structural and functional architectural facsimiles between the newborn and adult brain (Ball et al., 2014; Telford et al., 2017; Stoecklein et al., 2019). Hierarchical complexity (HC) is a new network measure characterizing the diversity of connectivity patterns found across hierarchically equivalent network nodes (i.e. nodes that have the same degree). Importantly, it distinguishes connectomes from different random null models where other common metrics fail, and is observed to reflect the hierarchical and functionally diverse capacities of the human brain (Smith et al., 2019), while not being a general feature of most other real-world networks (Smith, 2019b). Studying the degree hierarchy has also provided a tractable signature of brain network architecture in the adult connectome: four hierarchical tiers broadly comprise different categories of functional processing – cognitive, sensorimotor, integrative, and memory and emotion (Smith et al., 2019).

In this work, we aimed to establish whether hierarchical complexity observed in adults is already detectable in the newborn period, supporting the hypothesis that HC is an intrinsic property of human brain networks, arising from organizational principles that drive brain development. We also ask whether atypical brain development after preterm birth is reflected in HC differences, relative to term infants. To achieve these goals, we developed an approach to assess HC in the newborn period. This approach first introduces a systematic method to define the hierarchical structure of connectomes using group-aggregated degree distributions. We establish this in a cohort of neonates alongside a cohort of healthy adults for reference. We investigate resemblance in the connectome degree hierarchy between birth and adulthood. Finally, we explore the effect of preterm birth on HC in the newborn period.

## Material and methods

### Participants and data acquisition

For the present work, two datasets were used

### Theirworld Edinburgh Birth Cohort (TEBC)

Participants were recruited as part of a longitudinal study designed to investigate the effects of preterm birth on brain structure and long term outcome (Boardman et al., 2020). The study was conducted according to the principles of the Declaration of Helsinki, and ethical approval was obtained from the UK National Research Ethics Service. Parents provided written informed consent. A total of 136 neonates (77 preterm [with gestational age at birth *<* 32 weeks] and 59 term) underwent MRI at term equivalent age at the Edinburgh Imaging Facility: Royal Infirmary of Edinburgh, University of Edinburgh, UK. Details are provided in table 1. Of the preterm infants, 22 had bronchopulmonary dysplasia, 5 had necrotising enterocolitis and 3 required treatment for retinopathy of prematurity.

**Table 1:** Neonatal participant characteristics. The last column reports the uncorrected p-values of the group differences computed with t-test for normally distributed continous variables, Wilcoxon rank-sum test for non normally distributed continuous variables, and chi-squared test for categorical variables.

|  | **term (N=59)** | **preterm (N=77)** | **term vs. preterm** |
| --- | --- | --- | --- |
| PMA at birth (weeks) | 39.49 (36.42 - 42) | 29.48 (23.42 - 32) | $p = 1.98 \times 10^{-23}$ |
| Birth weight (grams) | 3484 (2556 - 4560) | 1278 (454 - 2100) | $p = 1.01 \times 10^{-52}$ |
| PMA at scan (weeks) | 41.84 (38.28 - 43.84) | 40.97 (38 - 44.56) | $p = 4.72 \times 10^{-5}$ |
| M:F ratio | 31:28 | 43:34 | $p = 0.8341$ |

PMA = Postmenstrual age, M = male, F = female.

A Siemens MAGNETOM Prisma 3 T MRI clinical scanner (Siemens Healthcare Erlangen, Germany) and 16-channel phased-array paediatric head coil were used to acquire 3D T2-weighted SPACE images (T2w) (voxel size = 1mm isotropic, TE = 409 ms and TR = 3200 ms) and axial dMRI data. Diffusion MRI images were acquired in two separate acquisitions to reduce the time needed to re-acquire any data lost to motion artifacts: the first acquisition consisted of 8 baseline volumes (b = 0 s/mm^2^ [b0]) and 64 volumes with b = 750 s/mm^2^, the second consisted of 8 b0, 3 volumes with b = 200 s/mm^2^, 6 volumes with b = 500 s/mm^2^ and 64 volumes with b = 2500 s/mm^2^. An optimal angular coverage for the sampling scheme was applied (Caruyer et al., 2013). In addition, an acquisition of 3 b0 volumes with an inverse phase encoding direction was performed. All dMRI images were acquired using single-shot spin-echo echo planar imaging (EPI) with 2-fold simultaneous multislice and 2-fold in-plane parallel imaging acceleration and 2 mm isotropic voxels; all three diffusion acquisitions had the same parameters (TR/TE 3400/78.0 ms). Images affected by motion artifacts were re-acquired multiple times as required; dMRI acquisitions were repeated if signal loss was seen in 3 or more volumes. Infants were fed and wrapped and allowed to sleep naturally in the scanner. Pulse oximetry, electrocardiography and temperature were monitored. Flexible earplugs and neonatal earmuffs (MiniMuffs, Natus) were used for acoustic protection. All scans were supervised by a doctor or nurse trained in neonatal resuscitation. Post acquisition, absolute and relative in-scanner motion were quantified by averaging voxel displacement across all voxels (computed as 3 translations and 3 rotations around the x, y and z axes) (Bastiani et al., 2019). Absolute displacement was computed with respect to the reference volume, while relative displacement was computed with respect to the previous volume. A summary measure for each subject was calculated as the average (absolute or relative) displacement across all volumes. There were no differences in either motion measure between the preterm and the term group (t-test *p >* 0.1).

### Human Connectome project (HCP)

We used the 100 Unrelated Subjects sample from the HCP 3T dataset, consisting of T1-weighted and dMRI data from 100 healthy subjects. Of these, 6 subjects were excluded because of known anatomical abnormalities^2^, resulting in a sample of 94 subjects (age range: 22-36 years; 44 male). T1-weighted data were acquired with 0.7 mm isotropic voxel size, TE = 2.14 ms and TR = 2400 ms. dMRI data were acquired with a 1.25 mm isotropic voxel size, TE = 89.5 ms and TR 5520 ms, with three shells with b = 1,000, 2,000 and 3,000 s/mm^2^, each shell with 90 diffusion weighted volumes and six non-weighted images (Essen et al., 2012).

#### Data preprocessing

##### Theirworld Edinburgh Birth Cohort

Diffusion MRI processing was performed as follows: for each subject the two dMRI acquisitions were first concatenated and then denoised using a Marchenko-Pastur-PCA-based algorithm (Veraart et al., 2016b); the eddy current, head movement and EPI geometric distortions were corrected using outlier replacement and slice-to-volume registration (Andersson et al., 2003; Andersson and Sotiropoulos, 2016; Andersson et al., 2016, 2017); bias field inhomogeneity correction was performed by calculating the bias field of the mean b0 volume and applying the correction to all the volumes (Tustison et al., 2010). The T2w images were processed using the minimal processing pipeline of the developing human connectome project (dHCP) to obtain the bias field corrected T2w, the brain masks, the tissue segmentation and the different tissue probability maps (Makropoulos et al., 2014, 2018). For the parcellation, the ten manually labelled subjects of the M-CRIB atlas (Alexander et al., 2017) were registered to the bias field corrected T2w using affine and symmetric normalization (SyN) (Avants et al., 2008), and then the registered labels of the ten atlases were merged using joint label fusion (Wang et al., 2013), resulting in a parcellation containing 84 regions of interest (ROIs). The 5 tissue type file needed to perform the tractography was generated by combining the tissue probability maps obtained from the dHCP pipeline with the subcortical structures derived from the parcellation process (https://git.ecdf.ed.ac.uk/jbrl/neonatal-5TT), an overview of the results can be seen in Figure 1a and Figure 1b. Finally, the mean b0 EPI volume of each subject was co-registered to their structural T2w volume using boundary-based registration (Greve and Fischl, 2009), then the inverse transformation was used to propagate ROIs label and the 5 tissue type file to dMRI space.

**Figure 1:**
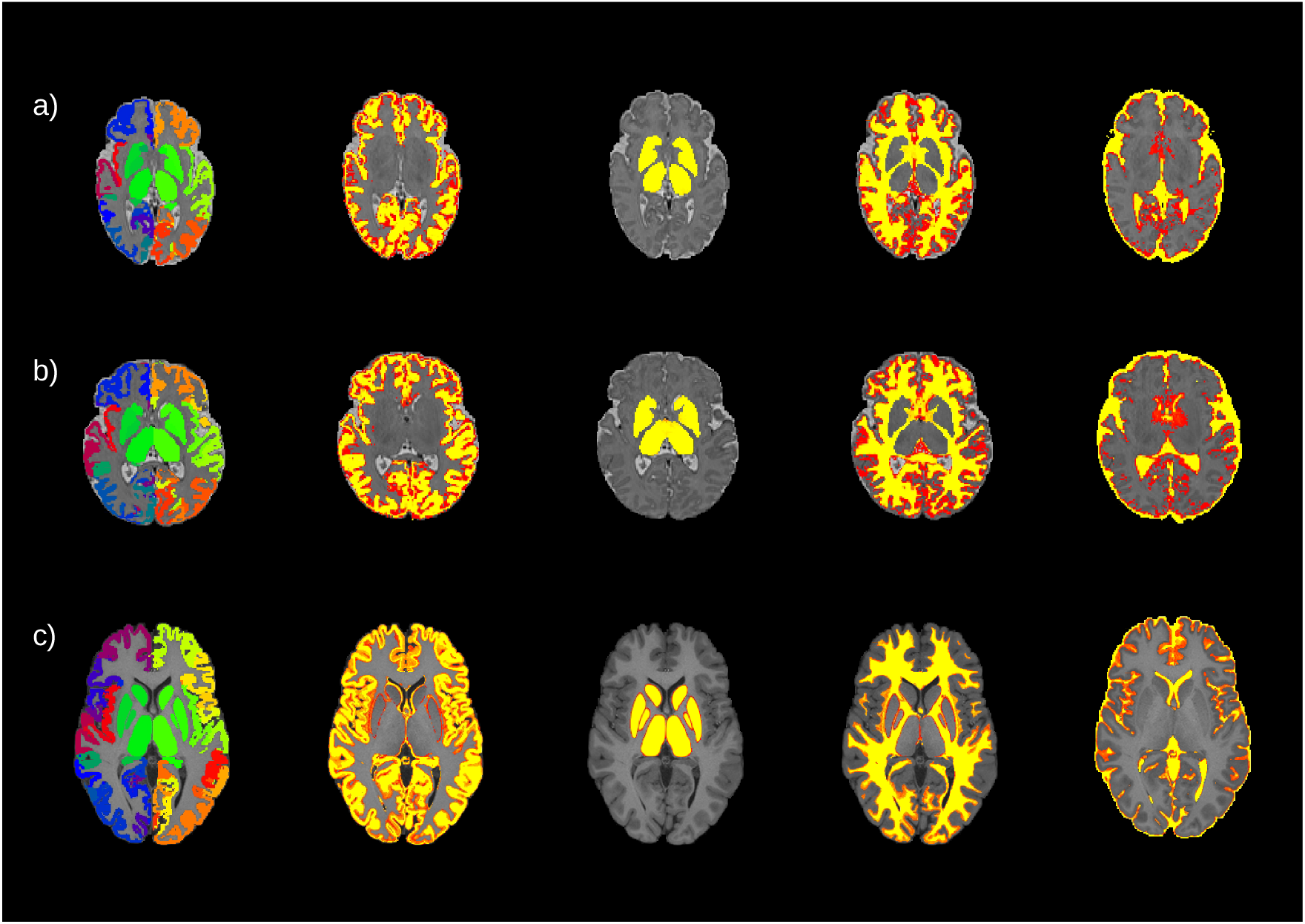
An example of the parcellation and the segmentation from three different subjects: a) a preterm born baby, b) a term born baby and c) an adult subject. From left to right, the parcellation and the four different tissue probability maps included in the 5 tissue type file: gray matter, subcortical gray matter, white matter and cerebrospinal fluid. For the neonates, the maps are overlaid onto the T2w volumes for the neonates and onto the T1w volume for the adult.

#### Human Connectome Project

The HCP dataset was already preprocessed, as described in (Glasser et al., 2013). Briefly, dMRI data were corrected for head motion and geometrical distortions arising from eddy currents and susceptibility artifacts (Sotiropoulos et al., 2013). Finally, the dMRI were aligned to the structural T1 image. The T1w was parcellated using the Desikan-Killany parcellation (Desikan et al., 2006), resulting in 84 ROIs. Using the T1w, the probability maps of the different tissues were obtained to create the 5 tissue type file (Zhang et al., 2001; Patenaude et al., 2011) (Figure 1c).

#### Tractography and network creation

The tractography was performed using constrained spherical deconvolution (CSD) (Tournier et al., 2007). For both datasets, a multi-tissue response function was calculated (Dhollander et al., 2016), the only difference is that for the neonatal cohort the FA threshold of the algorithm was reduced to 0.1. For each cohort, the average response functions were calculated. Then, the multi-tissue fiber orientation distribution (FOD) was calculated (Jeurissen et al., 2014) with the average response function using a ℒ _max_ = 8. For the HCP dataset three FODs were calculated, one for each tissue type WM, gray matter (GM) and cerebrospinal fluid (CSF); while for the TEBC only two (WM/CSF). Finally, a joint bias field correction and multi-tissue informed log-domain intensity normalisation on the FODs images was performed (Raffelt et al., 2018).

Tractography was then performed with the iFOD2 algorithm (Tournier et al., 2010) using anatomically-constrained tractography (Smith et al., 2012), generating 10 millions of streamlines with a cutoff of 0.05 (default), using backtrack (Smith et al., 2012) and a dynamic seeding (Smith et al., 2015b). To accommodate for the difference in brain size between neonates and adults, the length of the fibers was set based on previous literature; with a minimum length of 20 mm (Neher et al., 2017; Blesa et al., 2019) and a maximum of 200 mm for the neonatal data (Blesa et al., 2019) and of 250 mm for the HCP dataset (Smith et al., 2012). To be able to quantitatively assess the connectivity, SIFT2 was applied to the resulting tractograms (Smith et al., 2015b).

The connectivity matrix was constructed using a robust approach, a 2 mm radial search at the end of the streamline was performed to allow the tracts to reach the GM parcellation (Smith et al., 2015a; Yeh et al., 2019). Each connectivity matrix was multiplied by their *µ* coefficient obtained from the SIFT2 process, because the sum of the streamline weights needs to be proportional to units of fiber density for each subject (Smith et al., 2015b, 2020). Resulting in:

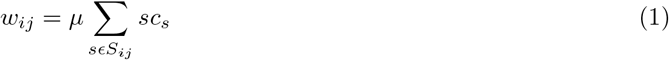

where *w*_*ij*_ is the total weight of the connection of the node *i* with the node *j, µ* is the coefficient obtained from SIFT2, *sc*_*s*_ is the SIFT2 weight of the streamline *s* and *s* ∈ *S*_*ij*_ represents all the streamlines connecting the nodes *i* and *j*.

As the connectivity matrices derived from probabilistic tractography are almost fully connected because of the presence of spurious connections, thresholding is often applied to obtain a sparser representation that is more likely to reflect the underlying network topology. We thresholded and binarized each individual connectivity matrices to obtain networks with a 30% density. This value is compatible with estimates from animal and human studies and was previously adopted for the study of human brain connectomes (Roberts et al., 2017; Buchanan et al., 2020) and for similarly generated networks (Roberts et al., 2016). To ensure that results were not biased to the selected threshold, we ran both the tier modelling and the main HC experiments across a range of thresholds (from 0.2 up to 0.4 in steps of 0.0005), see Supplementary material sections S1 and S2. Results were found to be consistent within this range.

#### Hierarchical complexity

The neighborhood of a network node is the set of all nodes with which that node shares links, and the number of the neighbors (and links) of a node is the nodal degree. The neighborhood degree sequence of the node is then the ordered sequence of degrees of the node’s neighbors, which is a particularly useful tool for studying organizational principles of networks (Smith, 2019b). The hierarchical complexity of a network involves computing the variability of neighborhood degree sequences of nodes of the same degree. This provides a measure of the diversity of connectivity patterns within the network degree hierarchy (Figure 2). Let *G* = (*V, E*) be a graph with nodes *V* = {1, …, *n*} and links *E* = {(*i,j*): *i,j* ∈ *V*}, and let *Κ* = {*k*_1_, *…, k*_*n*_} be the set of degrees of *G*, where *k*_*i*_ is the number edges adjacent to node *i*. Further, let Κ_*p*_ be the set of nodes of degree *p*. For neighborhood degree sequence 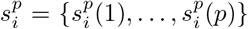 of node *i* of degree *p*, the hierarchical complexity is

**Figure 2:**
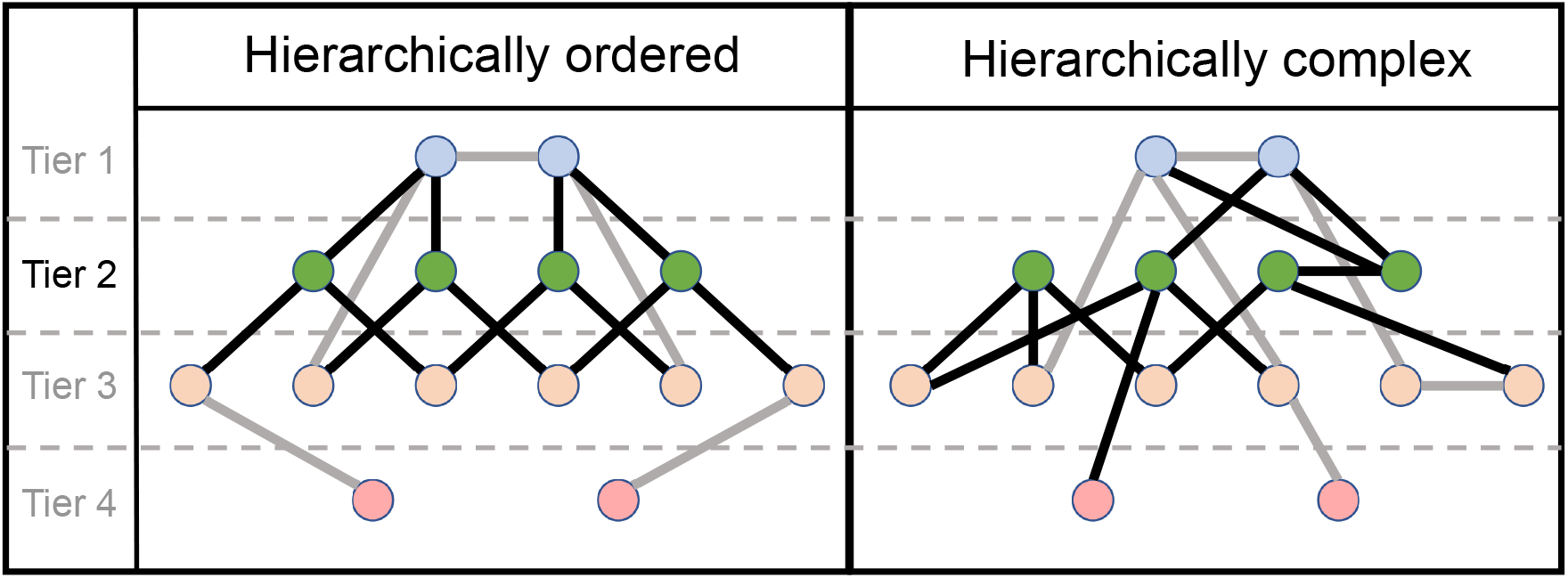
Illustration of ordered and complex hierarchical networks. Tiers are determined by the degree of nodes, with nodes with the largest degrees in the top tier. Tier 2 nodes and links are highlighted. In an ordered hierarchy, all nodes of similar degree are connected in a similar fashion, while a complex hierarchy is characterized by heterogeneous connectivity patterns across similar degree nodes.

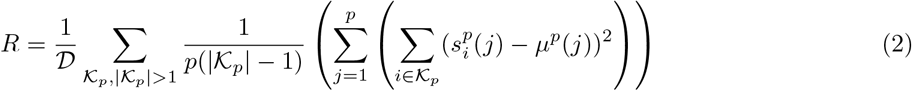

where 𝒟 is the number of distinct degrees in the network and *µ*^*p*^(*j*) is the mean of the *j*th entries of all *p* length neighborhood degree sequences (Smith and Escudero, 2017). To compute the hierarchical complexity within a hierarchical tier (defined as the set of nodes with a given range of degrees), we used degree specific hierarchical complexity by averaging hierarchical complexity over the range of degrees specific to the considered tier.

Many network analyses are conducted at nodal levels. However, in general, complexity is a system-level description of interacting components, and so, though we can provide a measure of hierarchical complexity across hierarchical levels, we are unable to provide a working definition of hierarchical complexity for single nodes. A tier level analysis, as we conduct here, is the most natural way of breaking down hierarchical complexity into finer detail.

#### Configuration models

To control for the differences in degree distribution between individual connectomes and the different populations (term and preterm born and adult), we used configuration models (Maslov and Sneppen, 2002). The configuration model fixes the number of links at each node of the null model by providing each node with a number of ‘stubs’, the number of which is the node’s degree in the original network. Then, pairs of stubs are randomly chosen across all nodes to establish new links. This process is repeated until there are no stubs remaining, meanwhile the process is started again anytime a link is created which either attaches a node to itself or attaches two nodes which already have an established link. This results in a random network that has the same degree distribution of the original network, whereas any organization is disrupted.

#### Hierarchical tiers

In addition to studying the hierarchical complexity of the whole network, we performed a more refined analysis by dividing the nodes of the network into different tiers, defined on the basis of nodal degree. Previous work (Smith et al., 2019) split each network into four tiers based on maximum degree magnitudes, where each tier comprised a rounded 25% of degrees. The first tier comprised nodes in the top 25% of degrees in the network, the second tier comprised of nodes with the next 25% of largest degrees, and so on. In our study, however, we wished to assess whether the degree distributions would reveal such tiers directly. To this end, we studied the group-aggregated degree distributions, i.e. the distribution of the pooled degrees of all nodes of all subjects within a group. We implemented an automated computational procedure based on Gaussian Mixture Modelling (GMM) to determine the tiers of the connectomes. Essentially, the best GMM was chosen and the theoretical components suggested by that model were taken as the tiers of the connectomes. Subsequently, thresholds between tiers were defined at the point where GMM component distribution functions intersected. See Supplementary material, section S1 for full details. Of note, we are not aware of any previous literature which has looked at group-aggregated degree distributions before, which is certainly partly why these different components have, until now, remained unnoticed.

Once tiers were defined, we implemented tier-based analysis on both the structural connectomes and their configuration models for comparison. To track the consistency of tiers across groups (preterm born, term born, and adult) we computed the number of times each node appeared in a given tier across participants. For each tier, Pearson’s correlation coefficients were then computed across these node proportions between preterm and term, preterm and adult and term and adult.

#### Hemispheric symmetry in network neighborhoods and common connections

As a post-hoc analysis, to better characterize the network topology and understand the results showed by the hierarchical complexity, we investigated the effect of cross-hemisphere neighborhood symmetry within tiers to probe deeper into the complex organization underlying the neonatal connectomes, following the insight that higher symmetry is associated with higher order and thus lower complexity (Smith, 2019b). We defined a measure of hemispheric symmetry based on the simple matching index between the sets of connections of a pair of homotopic regions, normalized with respect to the expected number of matching connections between two sets of connections selected independently at random. For full details see section S3 of Supplementary Material. We also studied the percentage of common and uncommon connections within tiers, following the hypothesis that adults have more well established network architecture and have more common connections within tiers than neonates. An ROI was defined as commonly connected to a given tier if it shared links with at least 80% of that tier’s ROIs. An ROI was defined as uncommonly connected to a given tier if it shared links with at most 20%, but not none, of that tier’s ROIs.

#### Characterizing connections within tiers

To further characterize the hierarchical tiers, we examined the types of connections within each tier. More specifically, we assigned connections to different categories according to whether they were 1) central-cortical, 2) intra-hemispheric cortico-cortical, 3) inter-hemispheric cortico-cortical, 4) central, 5) cortico-cerebellar, 6) central-cerebellar or 7) cerebellar. To obtain a group level solution, we assigned each region to a tier if it was included in that tier in at least two thirds of the population and we built a group connectome where we retained connections that were shared by at least two thirds of subjects in the group. In neonates, we also studied the distribution of connection lengths per tier and per group (where individual values were normalized by total brain volume).

#### Statistical analysis

Wilcoxon rank sum tests were carried out to assess the significance of the differences of distributions of network index values between the structural connectomes and configuration models. Unless otherwise specified, the Benjamini-Hochberg false detection rate procedure was applied on all reported *p*-values at the strict level of *q* = 0.05. The cut-off of the false detection rate in this study was 0.0222 (i.e. maximum acceptable *p*-value), which reflects the large proportion of differences found here. The effect sizes were also computed with Cohen’s *d*.

### Data availability

The hierarchical complexity code can be found in (Smith, 2019a). Reasonable requests for original image data will be considered through the BRAINS governance process (www.brainsimagebank.ac.uk) (Job et al., 2017).

## Results

In this study, our first aim was to define the tier organization of the neonatal connectome, and our second was to understand patterns of hierarchical complexity within these tiers both generally with respect to null models, and specifically in the context of preterm birth. In each case, due to the large differences expected in brain structure as well as in data acquisition, processing and parcellation of brain images between neonate and adult groups, we used the adult group as a qualitative reference for the neonatal group, while group-wise quantitative analysis was reserved for comparisons between term and preterm born neonates.

### Hierarchical tier organization

First, we inspected the existence of tiers in the structural connectomes of both term-born and preterm-born neonate groups and adults by considering their group-aggregated degree distributions (i.e. all degrees appearing in all connectomes of the group), Figure 3. This revealed distributions composed of several components identifiable from several distinct peaks with troughs in between. To quantitatively decompose these distributions into distinct components, we performed Gaussian Mixture Modelling (GMM). We varied the number of components in the modelling, from 2 up to 6, to find the best fit. The GMM models with best fit were identified by minimizing the Bayesian Information Criterion (BIC), which finds the best trade-off of high accuracy and low model complexity (i.e. avoiding overfitting). For each group the model with four components was found to minimize the BIC. To check the consistency of this with respect to network density threshold, we varied the threshold between 0.2 and 0.4 in steps of 0.0005. The 4 component GMM was consistently the best model across thresholds and the theoretical Gaussian components derived by the model identified the same underlying components of the connectome distributions across thresholds. Due to the overlapping elements of the distributions, we set the threshold between tiers at the point where the probability density functions of two adjacent components crossed each other, i.e. the point where the next component begins to have more likelihood of being the source of the node at that degree. For full details on all of these considerations the reader is referred to the Supplementary material, Section S1.

**Figure 3:**
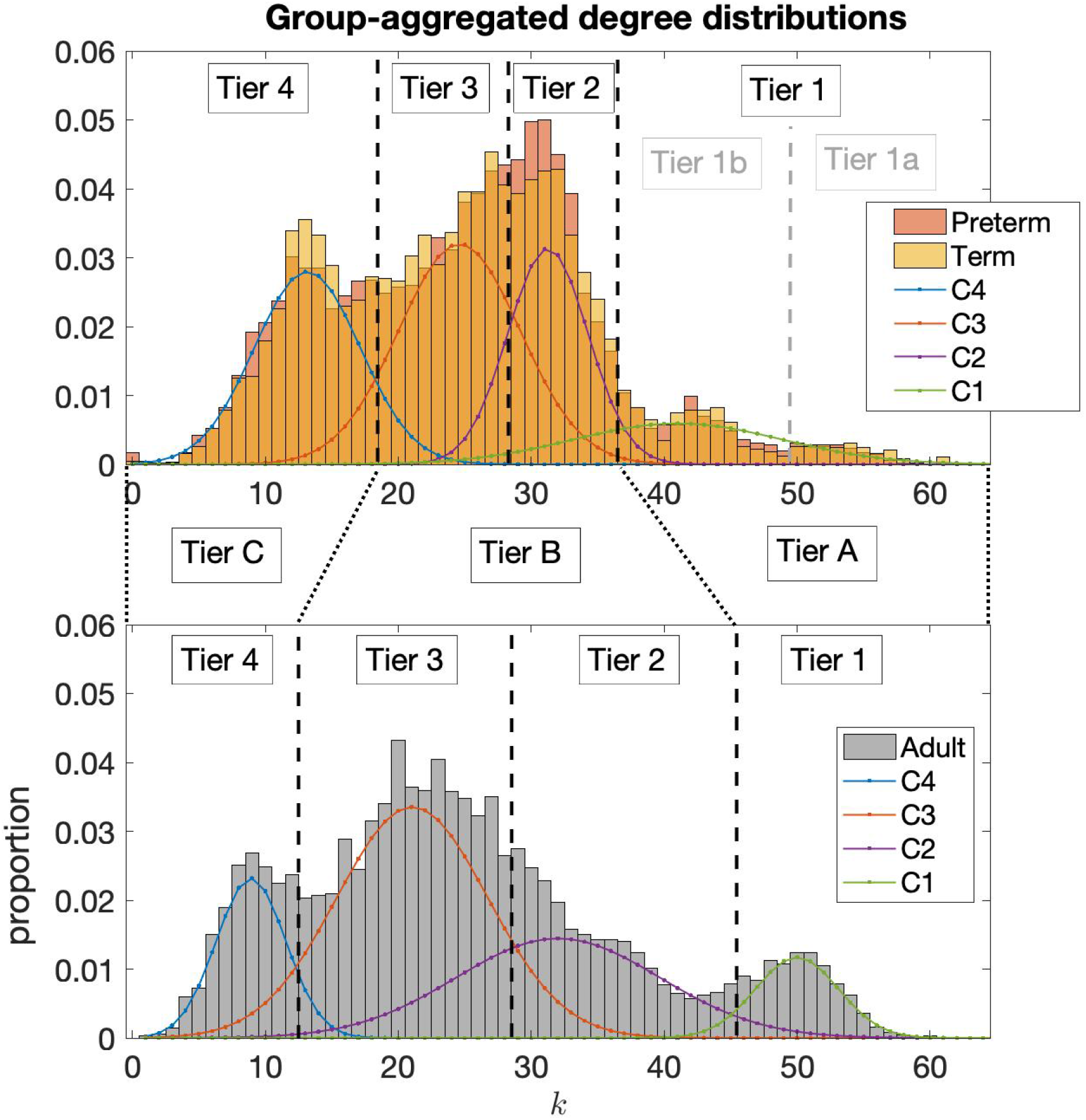
Aggregated degree distributions of neonatal groups, top, and the adult group, bottom. Four distinct peaks are noted in the degree distributions of neonatal connectomes and corresponding peaks are also seen in the adult connectomes. The four Gaussian Mixture Model components are shown as C4, C3, C2 and C1. These are taken as the natural tiers of the connectomes and black dotted lines indicate the thresholds between tiers. Greater consistency between neonates and adults is found by consolidating the tiers as indicated by Tier A, B and C.

Note, in the neonate group it was observed that the GMM failed to identify what was observable as two separate components at the right tail of the distribution, which we called Tier 1b and Tier 1a in Figure 3. Analysis of ROIs in these tiers showed that Tier 1a in neonates consisted solely of the thalamus, indicating the hyper-connectedness of this region in neonates. Indeed, consistency between neonates and adults was better achieved by consolidating Tier 1a and Tier 1b in neonates as Tier A, comparable to Tier 1 in adults. Correlations between ROI proportions in Tiers 2 and 3 between adults and neonates were significantly increased by consolidating these tiers as a single Tier B (from Pearson’s correlation coefficient of 0.5405 for tier 2 and 0.4949 for Tier 3 up to 0.6577 for the consolidated Tier B). This indicated that these components in adults and neonates were not directly comparable, while greater comparability was made possibly by combining them. Tier 4 ROIs did show high consistency between adults and neonates (Pearson’s correlation of 0.7505) and was simply relabelled as Tier C. See section S1 of Supplementary Material for full details.

We can see the 1 to 4 Tier distributions over the brain for each population in Figure 4. For this figure, an ROI was assigned to a tier if it was included in that tier in at least two thirds of the population. For the A-C Tier distribution see Supplementary Figure S6. The average number of nodes in each tier for the three groups of subjects is reported in Supplementary Table S2. The list of regions within each tier is provided as supplementary material (Table S3).

**Figure 4:**
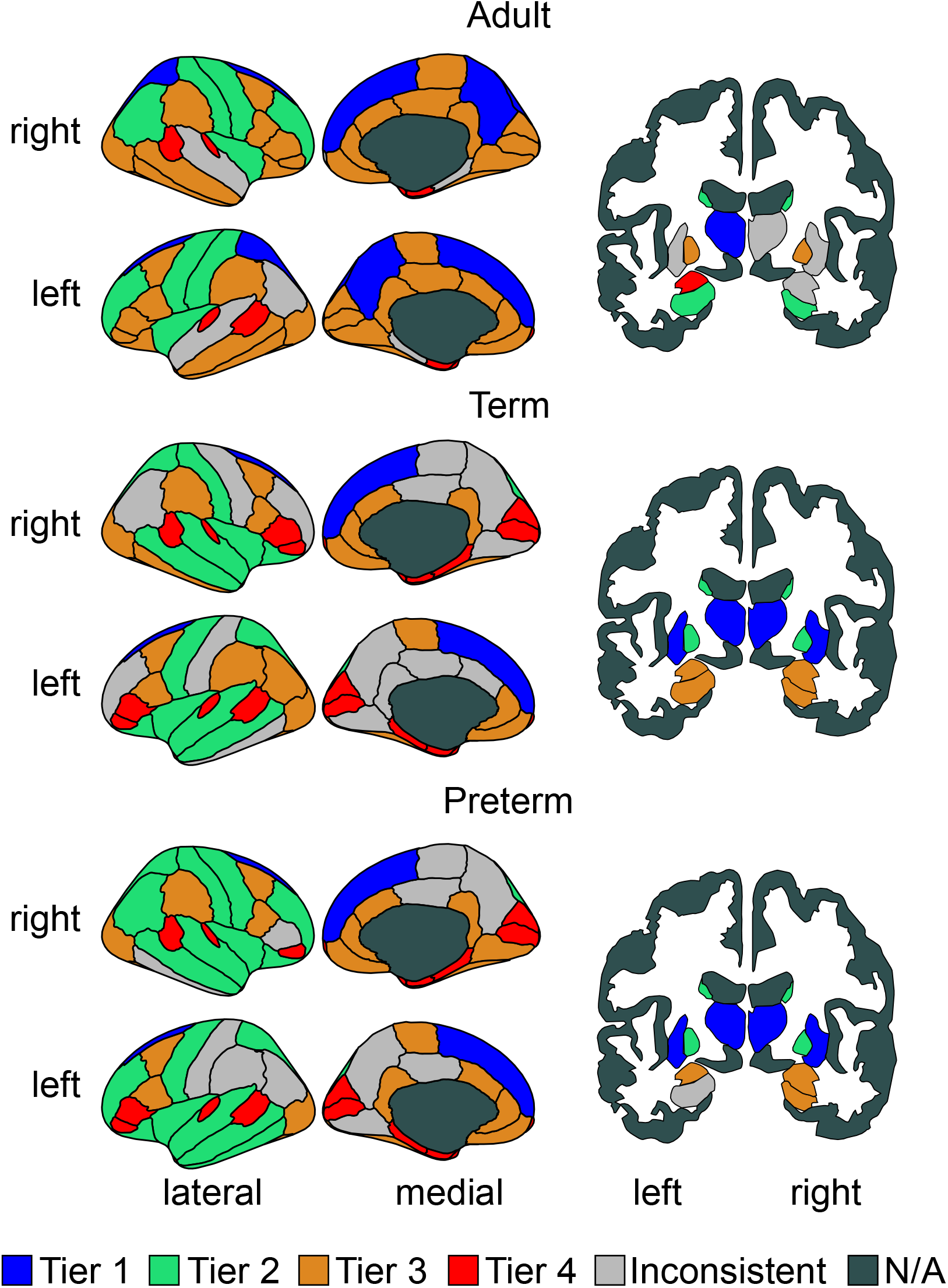
Cortical and sub-cortical representations colored by tier. N/A means non assigned. Due to the display plane used, two areas are not shown, the accumbens area, that was assigned (in both hemispheres) to Tier 4 in all three populations; and the cerebellum, that was assigned (in both hemispheres) to Tier 3 in all three populations.

### Hierarchical complexity

A very strong agreement was observed between the tiers of the neonate groups (Pearson’s correlations all *>* 0.95), whereas differences were noted between tiers in neonates and in adults, even after the consolidation of tiers described above (Supplementary Table S1).

We directly compared the global HC and HC within tiers for term and preterm born neonate groups using Wilcoxon rank sum tests while using adult HC as a qualitative reference, Figure 5. HC was significantly larger in term-born neonates than preterm-born neonates (*p* = 0.0148, *d* = 0.3946). Global HC of adult connectomes was observed to be much larger than both neonate groups, indicating the expected trend that the term group would exhibit characteristics between preterm and adults. As the preterm and term group differed in mean PMA at the time of scanning (Tab. 1), to test for potential confounding effects we measured Pearson’s correlation between age and global HC for the whole neonatal group and for the term and preterm group separately, and found no significant association (uncorrected *p >* 0.3). This is in agreement with previous results from Smith et al. (2019), where no association with age was found in adults. We also tested whether within the two groups HC was related to PMA at birth or birth weight, and none of the correlations was significant (*p >* 0.1). In addition, we tested the association between HC and absolute and relative in scanner motion, and also in this case we found no significant correlation (*p >* 0.06).

**Figure 5:**
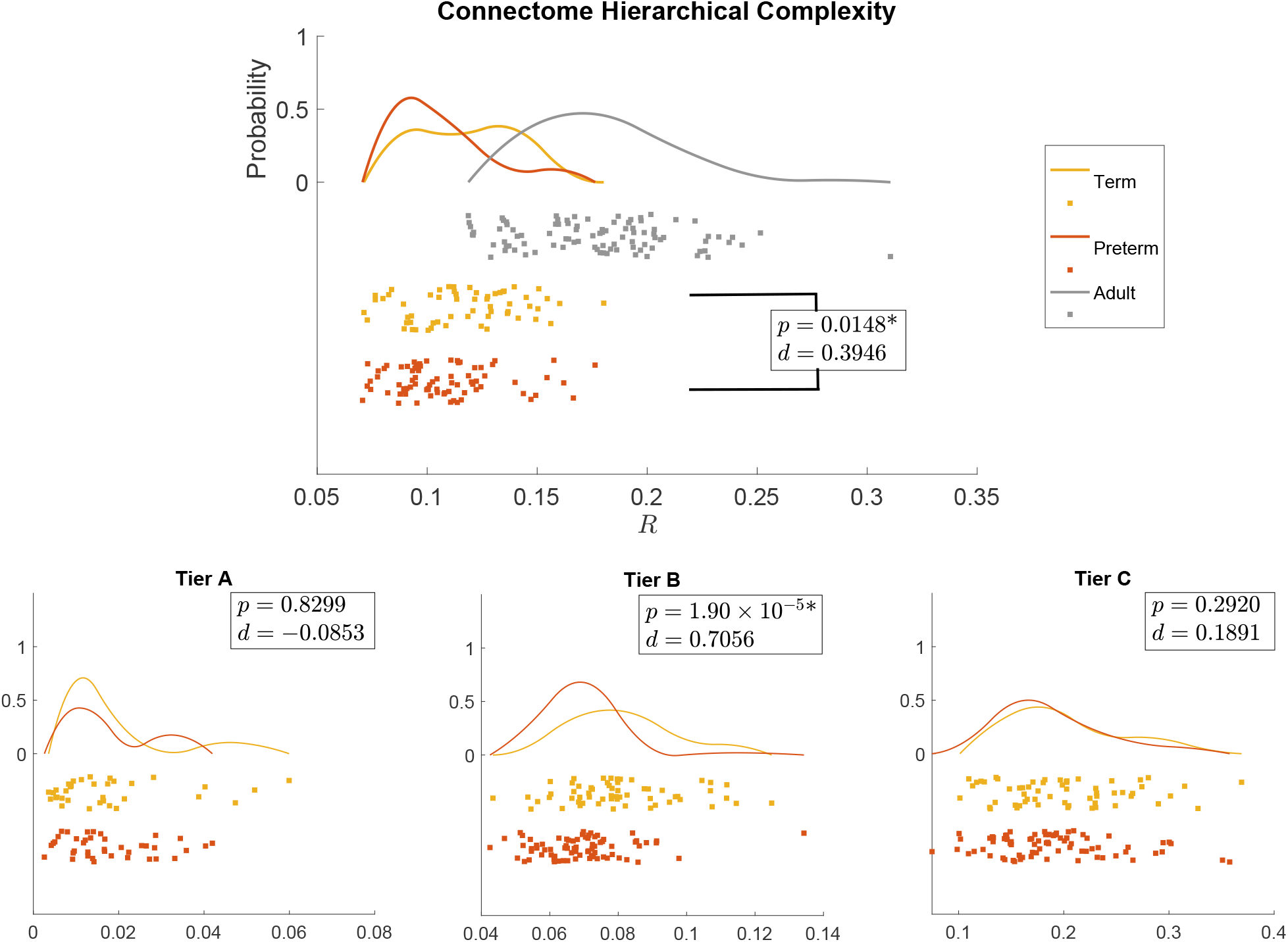
Distribution of the global hierarchical complexity (*R*) for the three populations as rain cloud plots (top) and hierarchical complexity of the three tiers in neonatal populations (bottom). Wilcoxon rank sum *p*-values and Cohen’s *d* values are shown for preterm vs term (all) and term vs adult (top). * denotes significant difference after FDR correction.

In the tier-based analysis, Tier B showed a corresponding significant difference in HC with a stronger effect size (*p* = 1.9 × 10^*-*5^, *d* = 0.7056), while no difference was evident in any other tier. Since global HC is a linear combination of the HC in each tier and no effect was found in Tiers A or C, we inferred that Tier B is likely the main cause of the global effect. To investigate this possibility we perfomed 2-way non-parametric analysis of variance for group × tier interaction. While both main effects were significant, the interaction term was not; this relationship requires further evaluation with a larger sample.

### Connectome Hierarchical Complexity

To get a general understanding of the trends of connectome complexity globally and in each tier, we compared each complexity value against those obtained for their corresponding configuration models (networks with the same degree distributions but randomized connections). We conducted this analysis for the adapted tier system to compare similar tiers across neonates and adults. The results are shown in Figure 6. The findings of global HC were confirmed in comparisons with configuration models with term-born connectomes having significantly greater HC than their configuration models, an effect which was not seen in preterm-born connectomes after False Discovery Rate (FDR) correction. An even larger significant difference was seen in adults.

**Figure 6:**
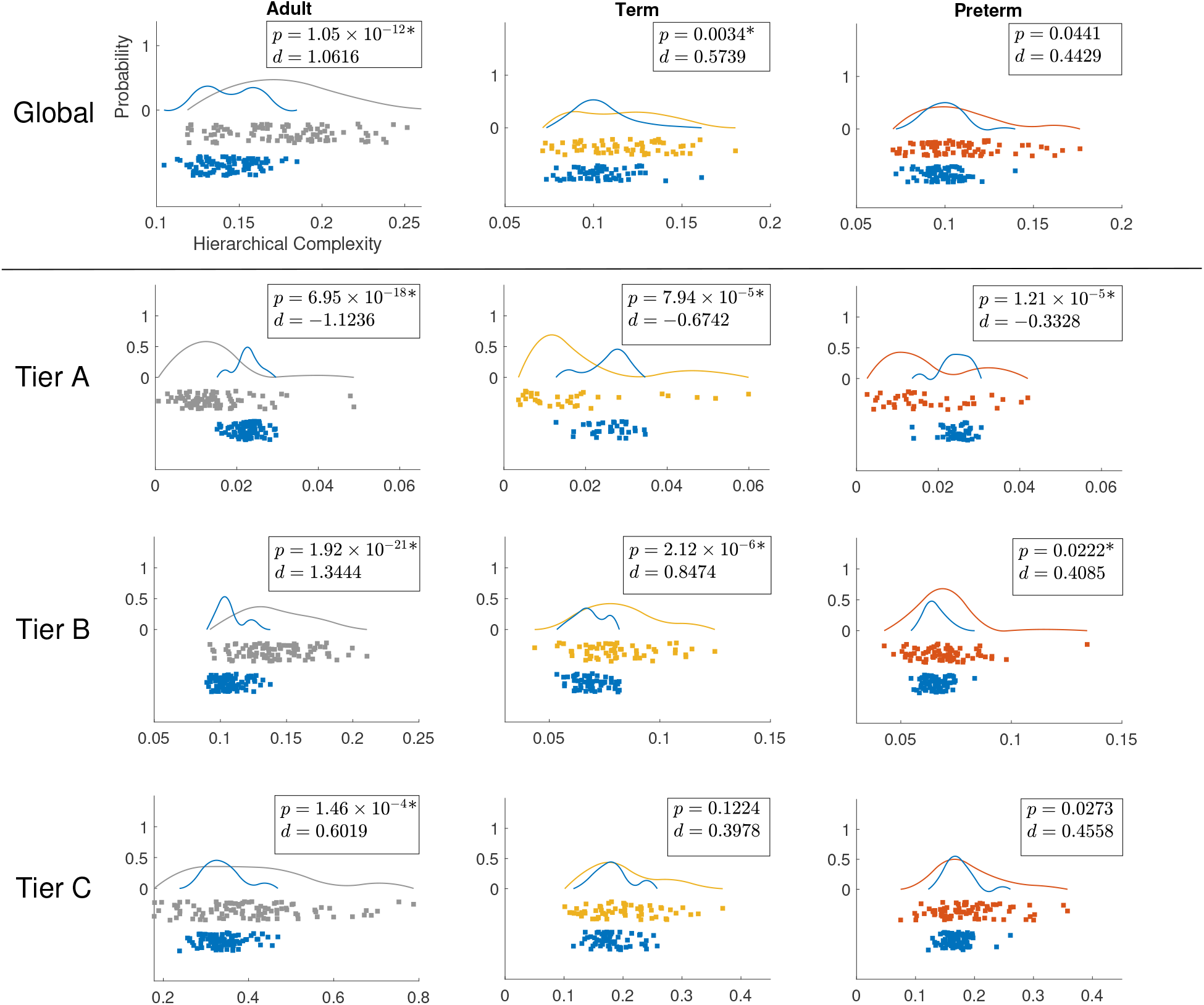
Distributions of hierarchical complexity globally (top) and for the different tiers of the three populations. Grey, yellow and orange colors represent values for adults, term born and preterm born neonates, respectively, while blue represents the values of hierarchical complexity for their corresponding configuration models. Wilcoxon rank sum *p*-values and Cohen’s *d* values are shown in top right corner of each plot. Axes as in top left plot. * denotes significant difference after FDR correction.

The result of greater complexity in Tier B of term-born neonates was also confirmed in comparisons with configuration models (Tier B in Figure 6). Tier B showed a clear increase of complexity compared to configuration models from preterm-to term-born babies. Interestingly there was also a clearly larger significant difference in adults. Surprisingly, Tier A exhibited lower HC compared to configuration models in all of the groups. This difference was evidently greater for adults compared to neonates. While the effect size was again larger in term-than in preterm-born neonates in Tier A, although with comparable *p*-values. Again, this suggested that the trend in term-born neonates moved away from preterm-born neonate connectomes towards adult connectomes. Since configuration models are the random null case, these results indicate that Tier A is more ordered than expected by random chance and becomes more ordered with maturation, while Tier B is more complex than random chance and becomes more complex with maturation.

#### Symmetry and common connections

We hypothesized that decreasing complexity in Tier A and increasing complexity in Tier B from neonates to adults reflected a systemic change whereby hub nodes create an ordered core structure integrating information from lower order nodes able to specialize into specific functional roles. To shed more light on the neurobiological contributions behind our results, we tested two possible contributors to the observed trends of HC: i) the hemispheric symmetry of connectivity patterns, and ii) commonality of ROIs connected to a given tier.Theoretically, hemispheric symmetry of connectivity patterns would influence HC because symmetric connectivity patterns would reduce HC and asymmetric connectivity patterns would increase HC (Smith, 2019b). Similarly, if the ROIs in one tier all tended to connect to the same other ROIs in the connectome, then this would decrease HC, and vice versa.

Table 3 reports the computed symmetry scores comparing the neighborhoods of homotopic brain regions, averaged by group and by tiers. We observed a consistent behavior across groups of subjects, with Tier A being the most symmetric, and decreasing symmetry from top to bottom tiers. Because it can be expected that nodes with greater degree will have a larger overlap of symmetric connections, just due to having many connections and a limited number of nodes with which to make those connections, we were cautious of interpreting these results as a general pattern. The results suggested that adults in general had less hemispherical symmetry across the connectome. From this, we concluded that hemispheric symmetry was not a strong contributing factor to HC, particularly increased symmetry in Tier A was not evident and not contributing to the observed increased order of Tier A nodes.

**Table 2:** Average degree by tier (mean ± standard deviation) and *p*-values of group differences from Wilcoxon rank-sum tests. * denotes significant difference after FDR correction.

| | average degree | | | $p$-value |
| --- | --- | --- | --- | --- |
|  | adult | term | preterm | term vs preterm |
| Tier A | 50.31±0.96 | 44.44±1.32 | 43.96±1.42 | 0.0429 |
| Tier B | 25.40±0.62 | 27.48±0.43 | 27.67±0.34 | 0.0104* |
| Tier C | 8.76±0.59 | 13.04±0.45 | 12.77±0.60 | 0.0033* |

**Table 3:**
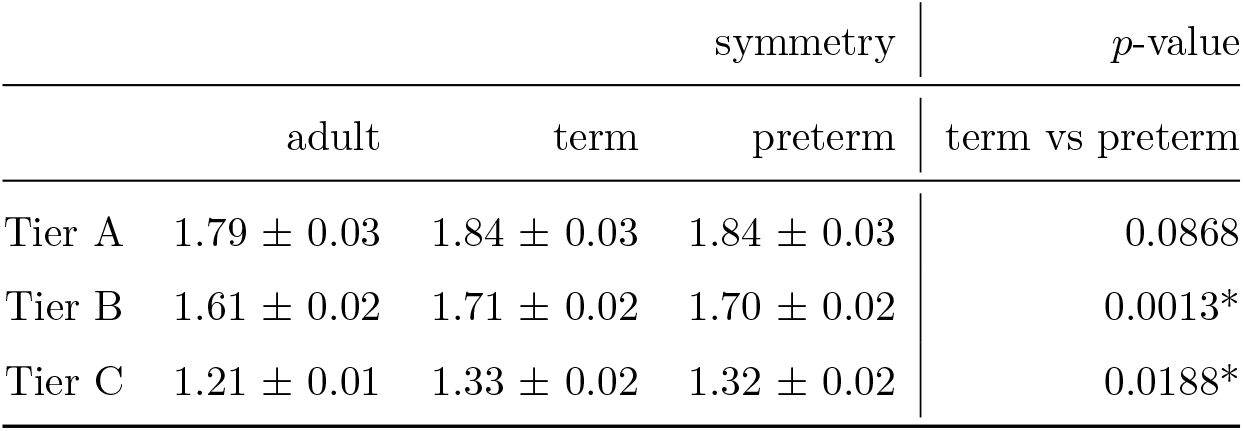
Symmetry scores by tier (mean ± standard deviation) and *p*-values of group differences from Wilcoxon rank-sum tests. * denotes significant difference after FDR correction.

Another possible contributing factor to the increased order seen in Tier A and increased complexity seen in Tier B would be if all Tier A ROIs tended to connect more to the same ROIs across the connectome while Tier B ROIs tended to exhibit less common connections in adults. To study this, we measured the number of common and uncommon connections made by nodes in each tier. See section S4 of Supplementary Material for full details. While no significant differences were found between term and preterm groups in any tier, Table 4, general trends between adult and neonate connectomes aligned with results of HC, with adults exhibiting more common connections than neonates in Tier A and more uncommon connections than neonates in Tier B, indicating that common and uncommon connections were likely a contributing factor to HC trends. This was further backed up by analysis of configuration models where neonates exhibited statistically less uncommon connections in Tier B to configuration models, while numbers of uncommon connections in Tier B for adults were not statistically distinguishable from configuration models, see supplementary figure S9.

**Table 4:** The *p*-values of group differences from Wilcoxon rank-sum tests between numbers of common and uncommon neighbors of each tier. * denotes significant difference after FDR correction.

|  | Tier A common | Tier B uncommon | Tier C uncommon |
| --- | --- | --- | --- |
| term vs preterm | 0.9439 | 0.2735 | 0.3948 |

#### Characterization of connections within tiers

The pie charts in Supplementary Figure S10 show the distribution of connection types for each group of subjects. In both neonate groups in Tier A the majority of connections are central-cortical (72/129 for term, 73/128 for preterm) and most of the central connections are in this tier (32/59 for term, 31/54 for preterm). Tier B has the highest proportion of intra-hemispheric cortico-cortical connections (352/812 for term, 368/829 for preterm) and most of the inter-hemispheric cortico-cortical connections (92/109 for term, 87/103 for preterm). Tier C has fewer connections but the pattern of cortical connections seems to mirror tier B. In comparison to neonates, adults have proportionally more intra-(34/128) and inter-hemispheric (45/128) cortico-cortical connections and fewer central (9/128) and central-cortical connections (34/128) in Tier A, and a slightly higher proportion of inter-hemispheric cortico-cortical connections in Tier B (134/682), while in Tier C there is a higher proportion of central-cortical connections (11/34) and a lower proportion of intra-hemispheric cortico-cortical connections (20/34). However, similarly to neonates, Tier B contains most of the cortico-cortical connections.

A chi-square test comparing connection type proportions in term vs. preterm neonates found no group difference. When considering connection type by tier in term vs. preterm neonates, we found a significant difference in the number of intra-hemispheric connections in tiers B and C, that were higher in the term group, and in the number of cortico-cerebellar connections in Tier B, higher in the preterm group (Bonferroni correction). The study of group aggregated connection lengths revealed that, compared to the other tiers, tier B had on average shorter connections in both neonates groups.

When comparing connection lengths between term and preterm groups with a Wilcoxon rank-sum test, preterm infants had longer connections in Tier A (*p* = 0.0001) and Tier B (*p* = 0.0011), while there was no difference in tier C (*p* = 0.9545), although overall the connection length distribution was similar in the two groups (Supplementary figure S11).

## Discussion

### Tier analysis

Empirical observations of group-aggregated degree distributions backed by GMM analysis revealed that the neonatal and adult connectomes are both comprised of four tiers; these findings were consistent across a range of network density thresholds. The original definition of hierarchical tiers was based on quartiles of the maximum degree, which were defined strictly for purposes of a more fine-grained analysis without any particular hypothesis of functionality (Smith et al., 2019). These current findings, combining empirical observation with statistical modelling, illuminate why that choice turned out to have biological relevance, with tiers matching quite closely different functional categories of nodes in adults.

Differences in tier composition were observed comparing adult and neonatal connectomes. The regions comprising Tiers 1a and 1b in the neonates were grouped together in the highest tier of the adult connectomes, while the regions comprising Tiers 2 and 3 in the neonates showed greater consistency with adult connectomes when these Tiers were combined. Correlations of ROI tier designation patterns were low for Tiers 2 and 3, but substantially increased when combining these Tiers (see Supplementary Table S1). This indicates a significant rearrangement of these medium degree ROIs in the developing brain.

Tier 1a in neonates was consistently composed solely of the thalamus. The thalamus forms a series of densely-connected processing loops with connectivity that covers large portions of the cortex and the hippocampus (Behrens et al., 2003; Grant et al., 2012; Aggleton et al., 2010). Among the subcortical regions, the thalamus and its connections are most strongly linked with supporting complex cognitive processes in adults, and its connections appear most strongly susceptible to ageing, partly via vascular risk (Cox et al., 2016, 2019b,a). In early life, prior work indicates that thalamo-cortical pathways are important for the regulation of area-specific differentiation of the developing brain (Boardman et al., 2006; Ball et al., 2013). The early wave of migrating neurons relay thamalo-cortical projections during late fetal and early preterm development, and the thalamic nuclei may exhibit asynchronous maturational trajectories: the posterior limb of the internal capsule (which include the posterior thalamic radiations) appears to develop earlier than the anterior limb (including anterior thalamic pathways (Hermoye et al., 2006; Dubois et al., 2014)). Tier 1 in adults, on the other hand, also contained the superior frontal gyrus and putamen (Tier 1b in neonates) as well as the precuneus (Tier 2 in neonates). The pattern of regional asynchrony in the developmental pattern of WM fiber organization and myelination may partly explain differences between neonatal and adult brain hierarchical organization. WM pathway growth and maturation spans a large time period, from fetal life through to late adolescence; during development frontal and temporal WM appears to show the most delayed developmental trajectories in fractional anisotropy (FA) (Deoni et al., 2012), although given that FA is likely to be differentially sensitive to the ‘local fiber architectural millieu’ in neonatal versus adult brains due to the differential presence of myelin, one should be cautious about directly attributing FA changes to any specific WM characteristic across life course. Given that subcortico-cortical and association fibres (cortico-cortical) especially in the frontal lobes exhibit later maturation (Young et al., 2017), it is likely that the relative shift up in degree of these regions represents more protracted course of hodological maturation in these areas.

Tier 2 in adults consistently contained the post central, precentral, rostral middle frontal gyri, and the insulate, caudate and hippocampus. Some of these can be broadly classified as lower order sensory processing regions; the poorer differentiation between HC tiers in neonates could reflect that a large amount of development of hierarchical organization occurs postnatally. For example, one could conjecture that the developmental period that ensues post-birth is the time during which there is hierarchical differentiation between sensory processing regions (Tier 2) and the heteromodal integration regions (Tier 3) that we see in the adult brain. Longitudinal research will be central to testing this hypothesis, and characterizing the changes in Tier membership that occur through childhood.

### Hierarchical complexity

The results on global complexity (Figure 5) indicate greater HC in the brains of infants born at term compared with brains of preterm infants at term equivalent age; and HC in adults was greater than in neonates. These observations indicate that HC is altered in association with early exposure to extrauterine life, and that HC of the structural connectome develops throughout childhood and adolescence until reaching the expected values of an adult healthy population. Comparing each population against their configuration model revealed that preterm complexity structure is not yet strongly distinguishable from random, whereas in term born babies, a more complex hierarchical structure is already present (Figure 6). As expected from previous results, this structure was clearly established in the adult dataset (Smith et al., 2019).

The tier-based analysis revealed that the global difference between term and preterm HC was due to the complexity evident in Tier B regions (figure 5), which has the largest number of ROIs in term and preterm infants (see Figure S6 in Supplementary Materials). At this age, the network segregation/integration balance is still undergoing many changes, including the transformation of the connectomic architecture from a relatively randomized configuration to a well-organized one (Cao et al., 2017).

When we compared the tiers against configuration models, we found that Tier A was less complex than its respective configuration model in all three populations while Tier B was more complex and no differences were found for Tier C, Figure 5. There were no differences in Tier A between term and preterm infants. Tier A corresponds to the hubs of the brain, and the lack of group difference in the neonatal period indicates that this “core network” is resilient to prematurity. Taken together, the findings are consistent with the hypothesis that term-born infants have a greater cerebral maturation than preterm-born infants, with topological properties that resemble more closely the properties of the adult connectome.

### Symmetry and common connections

Connectivity patterns in the adult brain were less symmetric than in neonates. This is consistent with recent findings reporting that most genetic effects on structural variation in the cortex are shared bilaterally and that asymmetry increases with age (Kong et al., 2018). It is reasonable to infer that the same is true of structural connectivity.

Surprisingly, we found that connectivity patterns in hub regions become more homogeneous with age and consistently more homogeneous than expected due to random chance. The fronto-parietal network has long been associated with higher cognitive processes such as general intelligence (Jung and Haier, 2007; Cox et al., 2019a). As such, it may be that postnatal development is important for establishing the emergent association fibre development that facilitates this networks’ support for higher order cognitive abilities.

The symmetry analysis and the comparison of common and uncommon connections lead us to conclude that hub regions are less symmetric in adults, while the connectivity patterns over all hub regions become more homogeneous.

### Characterization of connections within tiers

The analysis of connection types and lengths within tiers suggested that, overall, the increased complexity of tier B observed in the term and the adult group might be related to the presence of short-range cortico-cortical connections with highly variable connectivity patterns, while tier A is composed of a network of central-cortical regions with a more ordered structure, that is established early in development. When comparing connections between the term and the preterm group, the only differences were observed in the number of intra-hemispheric connections in tier B and C and the number of cortico-cerebellar connections in tier B. However it should be noted that in these analyses connections were aggregated by type, while HC is likely to be sensitive to the variability in how the regions are wired.

### Related work

Comparing neonatal structural connectivity studies is challenging, due to the lack of a standardized protocol for dMRI processing, parcellation or network analysis (for a review on the topic see Zhao et al. (2019b)). In the literature, an increased clustering coefficient, modularity, local and global efficiency and reduced characteristic shortest path length have been found in term-born infants compared against preterm-born infants scanned at birth (Tymofiyeva et al., 2013; Ball et al., 2014; Brown et al., 2014; van den Heuvel et al., 2015; Batalle et al., 2017). This translates in networks becoming more efficiently connected with development by achieving a trade-off between integration and segregation (Zhao et al., 2019b). Two studies comparing term-with preterm-born infants scanned at term equivalent age reported an increased small-worldness (Lee et al., 2019) and an increased clustering coefficient (Ball et al., 2014) in preterm infants. This suggests that the structural brain network after preterm birth is reorganized in maximizing integrated and segregated processing, implying resilience against prematurity associated pathology (Lee et al., 2019). Our findings also indicate that some aspects of connectome organization are resilient to preterm birth: there is at least a part of the network – the main hubs (Tier A) – that presents the same core structure at term in both term and preterm infants.

Although the general framework of brain circuits is dictated by genes and is in place by the time of birth (Keunen et al., 2017), the emerging brain networks are still immature, and chaotic unpredictable patterns of experiential signals from the environment together with altered physiology (inflammation, sub-optimal nutrition) can disrupt normal maturation (Short and Baram, 2019). Accumulating evidence from imaging studies supports the theory that preterm birth affects network maturation and brain structure: reduced WM and GM volumes, altered microstructure and atypical connectivity patterns (Boardman and Counsell, 2019), at global and local levels (e.g., in the thalamocortical system (Boardman et al., 2006; Ball et al., 2013)) and alterations in regions supporting neurocognitive and primary motor/sensory functions (Bouyssi-Kobar et al., 2018), all suggest delayed or atypical maturation associated with prematurity. Our results, showing a reduced HC in preterm-compared with term-born infants, corroborate this hypothesis and move towards providing a framework within which to observe and understand the developing brain as a complex hierarchical system.

Macro-scale connectivity gradients have been recently applied to study neonatal functional connectome organization, (Lariviére et al., 2020). The technique, which sorts brain regions along a continuous axis on the basis or their connectivity profile, revealed that although shortly after birth functional architecture is already set to allow for mediated mechanisms (e.g. thalamo-cortical integration), development prioritizes communication within the sensori-motor and visual systems, while higher order functional systems have an immature circuitry. This is in line with our finding that the hierarchical tiers in neonates lack the differentiation between sensory processing and association areas seen in adults, even if the overall network already presents a hierarchically complex organization.

A potential avenue for future research is to investigate whether nodal properties other than degree vary in association with network complexity, and to determine whether such properties enhance understanding of the hierarchical organization of the human brain during development.

## Conclusion

This study provides a new systems-level paradigm to understand the macro-scale developing brain. It is the first to consider the existence and implications of hierarchical tiers and their contingent connectivity patterns in the neonatal brain. We found that HC was greater in term-born neonates than in preterm infants. Natural tiers were discovered in the group-aggregated connectome degree distributions, with clear reconfigurations occurring between neonates and adults in high level and intermediate tiers. The tier-based analysis revealed that the difference in complexity between neonatal groups was greatest in neonatal Tier B, comprising regions involving sensorimotor processing and regions integrating high order cognitive and lower order sensorimotor processing. Comparisons with configuration models revealed a hierarchical structure where top tier hub regions were less complex (thus more ordered) than expected by random chance while Tier B regions were more complex than expected by random chance with statistical results indicating these patterns were dysmature in preterm-born neonates. The former result was at least partly due to common ROIs to which hub regions connected, with the superior frontal gyrus, putamen and precuneus joining the thalamus as hubs in adults. The complexity of tier B on the other hand indicated the beginnings of specialisation of multifarious cortical regions in neonates, with greater specialisation observed in term-born neonates. We have demonstrated the potential of this approach in a study of preterm birth, but these concepts can be applied in more general settings to understand the neural bases of cognition in health and disease. A natural extension of this work would be analysis of the developmental trajectory of HC across childhood and adolescence, and its variability in older age. This would allow the investigation of deviations from normal progression associated with cognitive impairment, and any brain disorder in early or later life that is characterized by alterations in network topology and global connectivity patterns.

## Funding

This work was funded by Theirworld (www.theirworld.org) and by Health Data Research UK (MRC ref Mr/S004122/1), which is funded by the UK Medical Research Council, Engineering and Physical Sciences Research Council, Economic and Social Research Council, National Institute for Health Research (England), Chief Scientist Office of the Scottish Government Health and Social Care Directorates, Health and Social Care Research and Development Division (Welsh Government), Public Health Agency (Northern Ireland), British Heart Foundation and Wellcome. MJT was supported by NHS Lothian Research and Development Office. Part of the work was undertaken in the MRC Centre for Reproductive Health, which is funded by MRC Centre Grant (MRC G1002033). SRC was supported by the MRC (MR/R024065/1) and National Institutes of Health (R01AG054628).

## Acknowledgments

Adult datasets were provided by the Human Connectome Project, WU-Minn Consortium (Principal Investigators: David Van Essen and Kamil Ugurbil; 1U54MH091657) funded by the 16 NIH Institutes and Centers that support the NIH Blueprint for Neuroscience Research; and by the McDonnell Center for Systems Neuroscience at Washington University. Individual parcellated templates and structural MRI images from the M-CRIB atlas (Alexander et al., 2017) were supplied by the Murdoch Children’s Research Institute. Neonatal participants were scanned in the University of Edinburgh Imaging Research MRI Facility at the Royal Infirmary of Edinburgh which was established with funding from The Wellcome Trust, Dunhill Medical Trust, Edinburgh and Lothians Research Foundation, Theirworld, The Muir Maxwell Trust and many other sources. We are grateful to the families who consented to take part in the study and to all the University’s imaging research staff for providing the infant scanning.

## Notes

### Competing Interest Statement

The authors have declared no competing interest.

