## Supplementary material for "Hierarchical complexity of the macro-scale neonatal brain"

---

*Keywords:* Hierarchical Complexity, Newborn, Developing Brain, Network Analysis, Structural Connectome, dMRI

---

### S1. Designation of tiers

The group-aggregated degree distributions (Figure 3 in main text) were clearly not unimodal distributions, exhibiting several noticeable distinct peaks suggesting they were composed of several components.

To confirm the validity of this observation, we performed Gaussian Mixture Modelling (GMM), implementing a standard linear regression fit (function *glmfit* in Matlab). GMMs were run 1000 times for models with 2, 3, 4, 5 and 6 components. For example, a model with four components is the aggregated distribution from four Gaussian distributions with different means, $\mu_1, \mu_2, \mu_3, \mu_4$ and variances $\sigma_1, \sigma_2, \sigma_3, \sigma_4$. The modelling also provides the proportions of each of these distributions, $w_1, w_2, w_3, w_4$ which determines contribution of a given distribution to the mixture. This gives the probability density function (pdf) of

$$p(x|\lambda) = \sum_{i=1}^{4} = w_i g(x|\mu_i, \sigma_i), \tag{1}$$

where each $g$ is the pdf of a Gaussian distribution with the mean and variance in the conditional argument, and $\lambda = \{w_i, \mu_i, \sigma_i\}_{i=1}^{4}$ is the collection of all parameters. For full details see e.g. Reynolds (2009).

Each modelling iteration was tested for fit against the real data using the Bayesian Information Criterion (BIC), which is lowest for models achieving a trade-off between high accuracy and low complexity, aiding to avoid selection of models which overfit the data. From a set of models, the best model is the one achieving

---

[1]These authors contributed equally to the work.

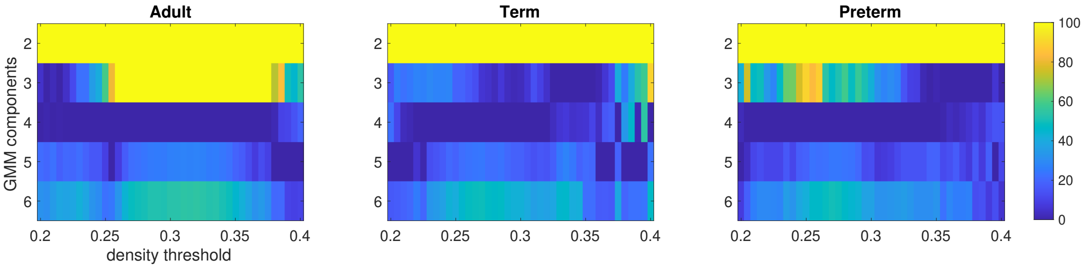


Figure S1: Average Bayesian Information Criterion values (relative to minimum across component number) for Gaussian mixture models applied to a range of density thresholds of the structural connectomes. The colour axis is cut off at 100 (bright yellow) to make comparisons easier between the more likely models. For each density, the minimum (dark blue) is the best model.

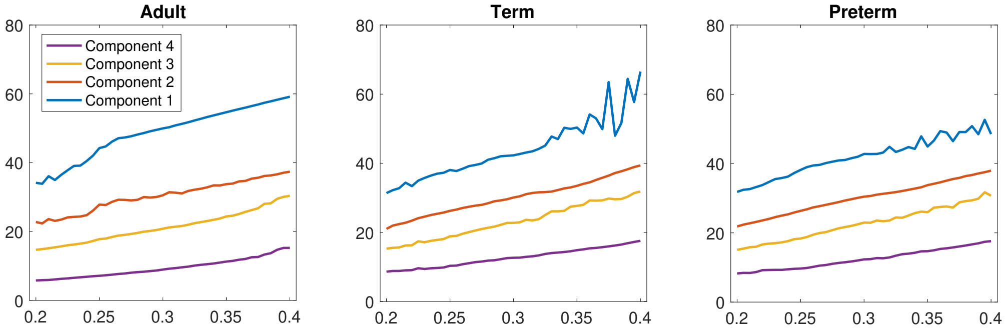


Figure S2: Average over 1000 iterations of the means of 4-component Gaussian mixture models (y-axis) applied to a range of density thresholds of the structural connectomes (x-axis).

smallest BIC. Fig S1 shows results for connectomes thresholded at different densities, from 0.2 up to 0.4 in steps of 0.0005. Those shown are BIC scores minus the minimum found over all models (2,3,4,5, and 6 component models) at that density, so that the minimum model at a given density has a value of 0. Results are averaged over 1000 iterations for each model. It is clear that the adult group was best represented by a 4 component model over almost all densities, while neonates were best represented mostly by 4 components, but also by 3 components at larger densities.

The means of the Gaussian components in the selected 4-component models are plotted– as averages over 100 iterations– in Fig S2. With increasing density (equivalently, number of links), the values for each component drifted upwards which was entirely expected since increasing density necessarily increases the average degree. Otherwise it was evident that the components related to the same underlying phenomena in the networks across thresholds.

To define tiers within adult and neonate groups, we used the 4-component GMM achieving minimum BIC score (out of the 1000 iterations) for each group. Note, to get a consistent tier definition for term and preterm born neonates, these groups were aggregated. From these models, we automatically set the threshold between tiers at the point where the probability density functions of two adjacent components crossed each other, i.e. the point where the next component began to have more likelihood of being the source of the node at that point, see Fig. 3 in the main text for details. Therefore, for neonates, Tier 1 was defined from the maximum degree down to a degree of 37, Tier 2 from 36 down to 29, Tier 3 from 28 down to 19 and Tier 4 from 18 down to 0. For adults, Tier 1 was defined from the maximum degree to a degree of 46, Tier 2 from 45 down to 29, Tier 3 from 28 down to 13 and Tier 1 from 12 down to 1.

For neonates it was notable that the GMM model was unable to pick up on the two smaller distribution components observable at the right-end of the distribution, instead treating them as a single distribution with a very large standard deviation. In our initial analysis, we kept these two observable components separate to study this further.

Next, for the best fit 4-component models, we inspected the consistency of the makeup of the ROIs in each component between groups. For each ROI and each group we counted how many times an ROI was assigned to each tier and divided these numbers by the group size. This provided the proportion of times an ROI was in each tier for that group. We then computed Pearson's correlation coefficients of these proportions over all ROIs between the groups, Table S1. Correlations between neonate groups were extremely high, with all tiers being above 0.95, indicating a strong alignment between the hierarchical structure of term and preterm neonate connectomes, see Fig S3. Correlations between neonate groups and adults were much more variable. While Tier 1 and Tier 4 showed moderately strong correlations, Tiers 2 and 3 had only medium correlations. However, on combining Tiers 2 and 3 together, the correlations significantly increased, Table S1, right. To make the analysis of more relevance for the adult and neonate groups, we thus designated an adapted tier system. We defined i) Tier A as Tier 1, ii) Tier B as the combination of Tiers 2 and 3, and iii) Tier C as Tier 4, in both neonates and adults. This is also reflected in the relative tier proportions between adults and term-born neonates using both the original 4-tier system, Fig S4, and the adapted 3-tier system, Fig S5. Figure S6 shows the spatial distribution of the tiers and table S2 the average number of nodes by tier within groups.

Table S1: Pearson correlation coefficients of ROI tier proportions between groups.

| | Tier 1 | Tier 2 | Tier 3 | Tier 4 | Tier A | Tier B | Tier C |
|---|---|---|---|---|---|---|---|
| preterm vs. term | 0.9856 | 0.9517 | 0.9554 | 0.9953 | **0.9856** | **0.9910** | **0.9865** |
| neonates vs. adult | 0.7241 | 0.5804 | 0.5012 | 0.7606 | **0.7241** | **0.6691** | **0.7606** |

Table S2: Average number of nodes by tier in adult and neonatal connectomes.

| | adult | term | preterm |
|---|---|---|---|
| Tier A | 9.22 | 5.92 | 5.74 |
| Tier B | 56.88 | 56.03 | 56.53 |
| Tier C | 17.89 | 22.05 | 21.72 |

### S2. Consistency of results across different network thresholds

Choosing a threshold for brain networks always contains some element of arbitrariness in selection. To ensure that the results of hierarchical complexity obtained in our study were stable to thresholding and not some artefact of random fluctuations, we ran the global and tier-based HC analysis on connectomes thresholded from 0.2 up to 0.4 in steps of 0.0005 (similarly as used for testing the reliability of the GMM modelling to threshold variations). The results– $p$-values of Wilcoxon rank sum tests between term and preterm born neonate groups– are shown in Fig S7. We see that the values achieved at the chosen threshold of 0.3 in our main study are entirely consistent with results obtained across this range of 401 thresholds, confirming the effect noticed is robust to threshold variations.

### S3. Hemispheric symmetry

We hypothesized that hemispheric symmetry contributed to hierarchical complexity, since ROIs with the same connectivity patterns should contribute to low complexity and vice versa. We wanted to know if hemispheric symmetry played a role in the trends of hierarchical complexity witnessed in the neonatal and adult groups. We defined a measure of hemispheric symmetry based on the simple matching index between the sets of connections of a pair of homotopic regions. In an adjacency matrix, each region is represented by a binary vector (a row or column of the matrix), where a 1 indicates the presence of a connection and 0 the absence of a connection. We compared each region with its symmetric counterpart, taking care of matching within and between hemispheric connections from each side (e.g. connections within the left hemisphere with connections withing the right hemisphere, left-to-right connections with right-to-left connections, and so on). For two binary vectors $x$ and $y$ of the same size $n$, the simple matching index is

$$SMI(x, y) = \frac{M11 + M00}{M11 + M00 + M01 + M10} = \frac{M11 + M00}{n} \qquad (2)$$

where $M11$ is the number of matching 1s in the two vectors, $M00$ is the number of matching 0s, $M10$ is the number of times there is a 1 in $x$ and a 0 in $y$ and $M01$ is the number of times there is a 0 in $x$ and a 1 in $y$. Now, we define the hemispherically mirrored adjacency matrix $A'$ as that obtained by switching rows and columns of all left and right pairs of ROIs. Taking the first vector $x_i$ as the row of $A$ corresponding

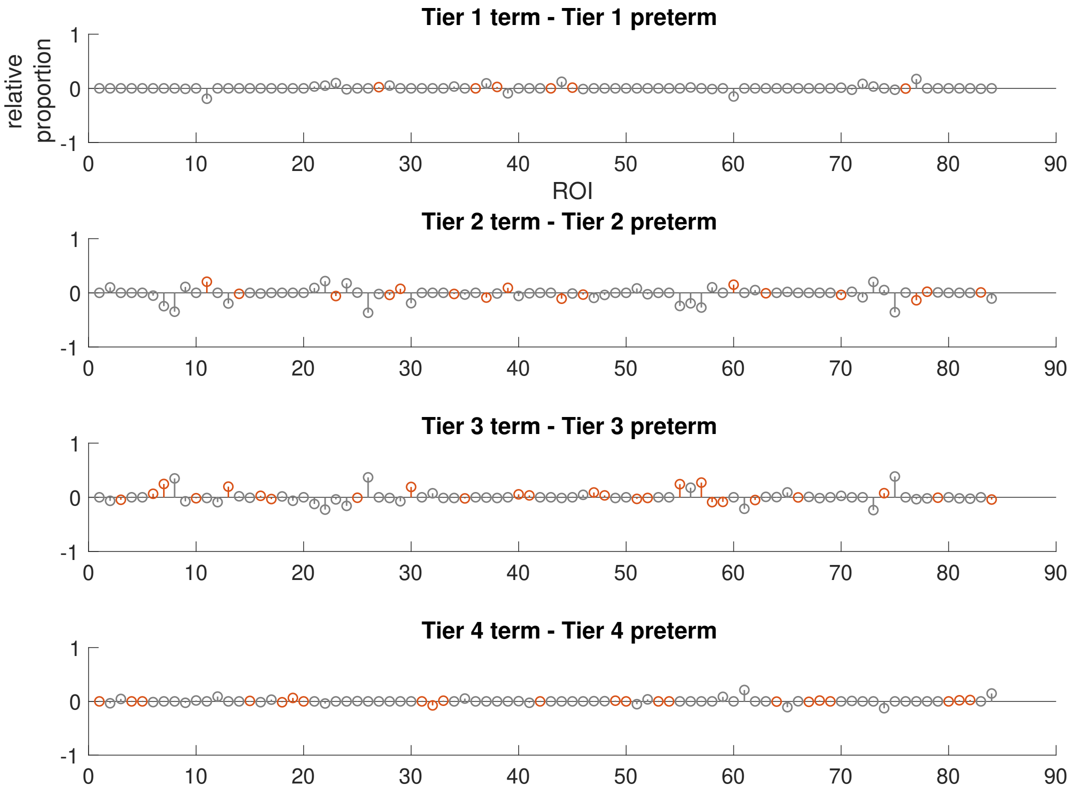


Figure S3: Difference in tier designation proportions of ROIs between term-born and preterm-born neonates in the four tier scheme. A value of 1 indicates the ROI is in that tier for all participants in the first named group in the plot title, while being in that tier for none of the second named group, and vice versa for -1. A value of 0 indicates the ROI is in that tier for the same proportion of participants in both groups. Orange nodes indicate the nodes in the relevant tier for the term group.

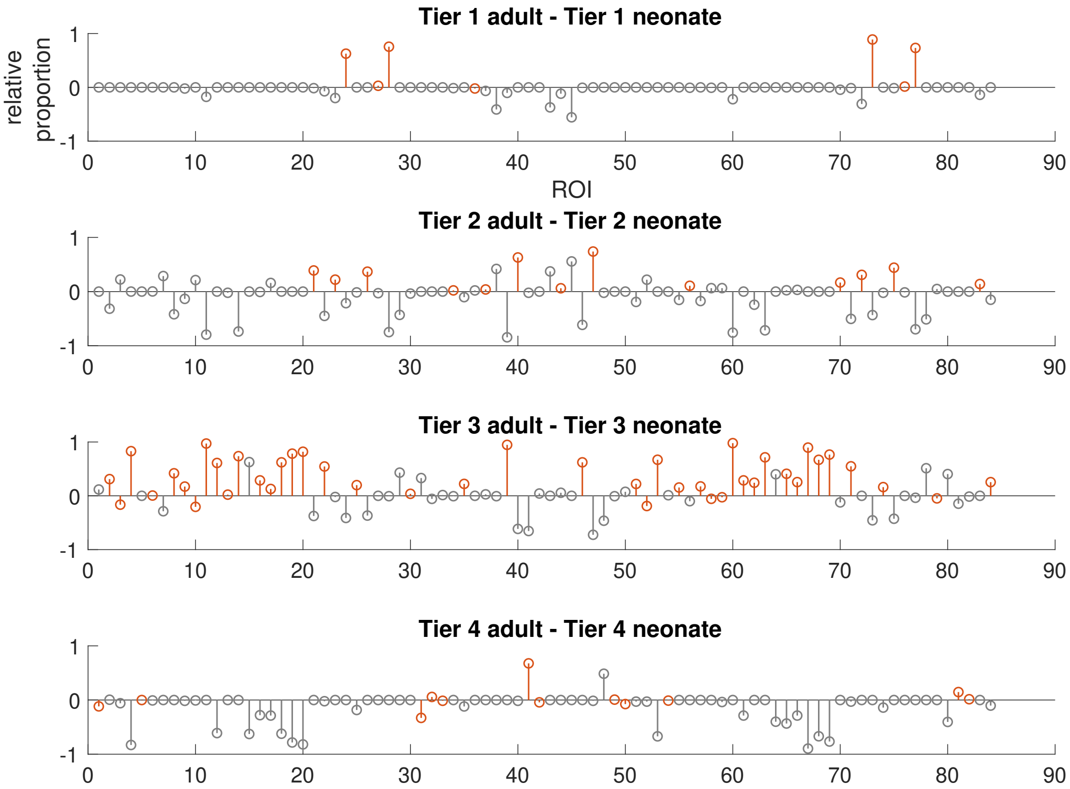


Figure S4: Difference in tier designation proportions of ROIs between adults and term-born neonates, directly comparing the tiers in the four tier scheme. A value of 1 indicates the ROI is in that tier for all participants in the first named group in the plot title, while being in that tier for none of the second named group, and vice versa for -1. A value of 0 indicates the ROI is in that tier for the same proportion of participants in both groups. Orange nodes indicate the nodes in the relevant tier for the adult group.

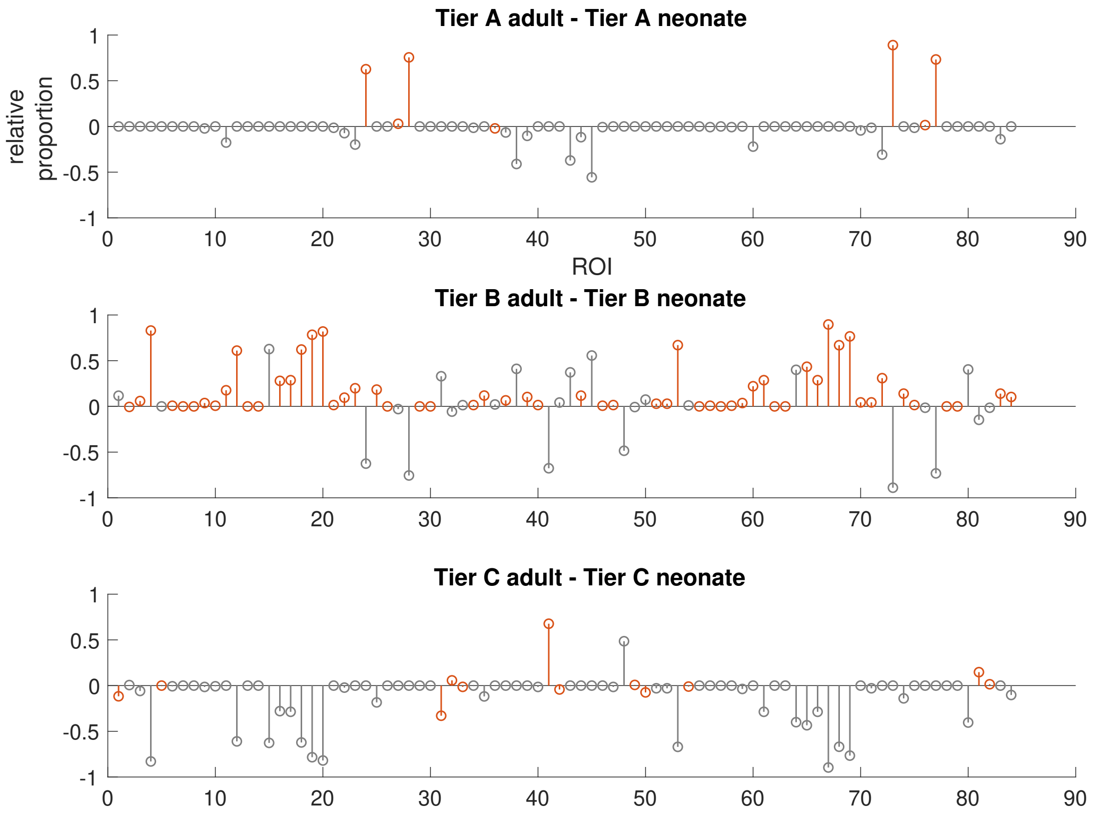


Figure S5: Difference in tier designation proportions of ROIs between adults and term-born neonates, comparing the adapted tiers A,B and C. A value of 1 indicates the ROI is in that tier for all participants in the first named group in the plot title, while being in that tier for none of the second named group, and vice versa for -1. A value of 0 indicates the ROI is in that tier for the same proportion of participants in both groups. Orange nodes indicate the nodes in the relevant tier for the adult group.

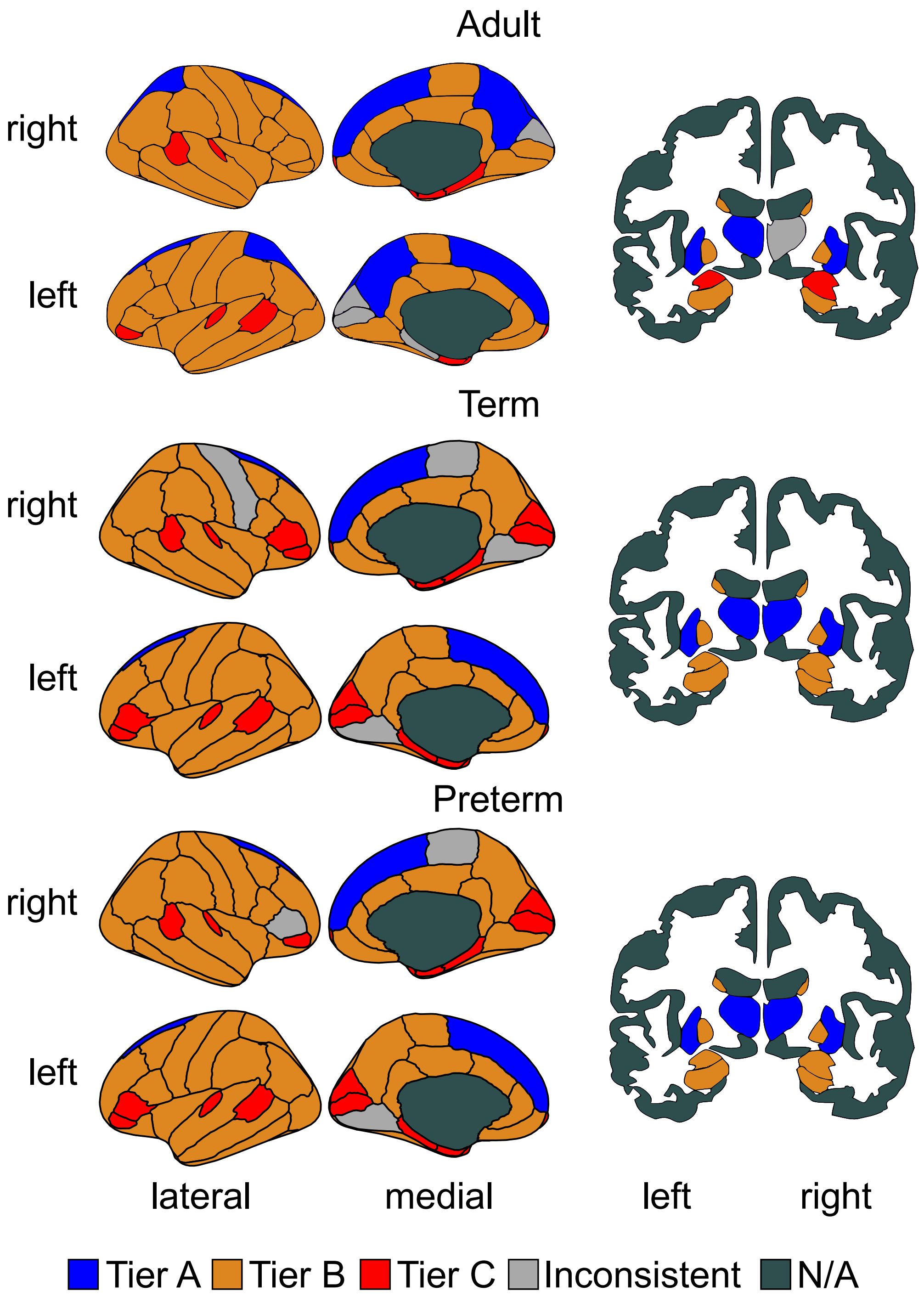


Figure S6: Cortical and sub-cortical representations colored by tier. N/A means non assigned.

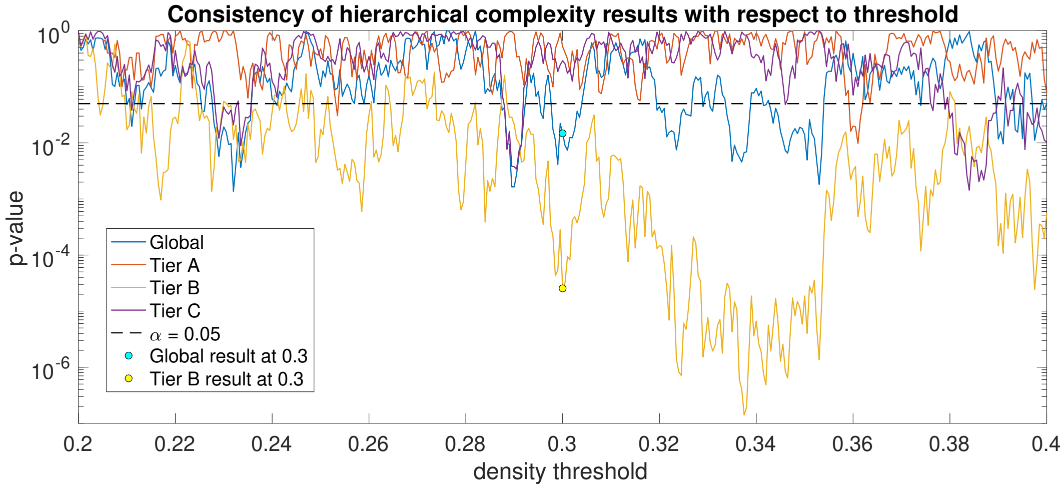


Figure S7: Results ($p$-values of Wilcoxon rank sum tests) of global and tier-based hierarchical complexity between term and preterm born neonate connectomes across different network density thresholds. Results for the threshold used in the main study (0.3) are highlighted.

to ROI $i$, and the second vector $x_{i'}$ as the row of $A'$ corresponding to ROI $i$'s symmetric counterpart, the hemispheric symmetry of a single ROI is defined as

$$HS(x_i, x_{i'}) = \frac{\sum_{j=1}^{n} \delta(x_i(j), x_{i'}(j))}{n} \tag{3}$$

where $\delta$ is the Kronecker delta function, being 1 if the two inputs are equal and 0 otherwise, $x_i(j)$ is the $j$th element of vector $x_i$ and $n$ is the maximum number of possible connections. We normalized our results with respect to the expected number of matching connections between two sets of connections selected independently at random. Let $V$ be the set of all possible connections and define $S_i$ as the set of connections of ROI $i$ with size $|S_i| = k_i$ and $S_{i'}$ as the set of connections of ROI $i'$ with size $|S_i| = k_{i'}$. Then

$$\begin{aligned} E[HS(x_i, x_{i'})] &= \frac{E[|S_i \cap S_{i'}| + |V \setminus S_i \cap V \setminus S_{i'}|]}{n} \\ &= \frac{k_i k_{i'}/n + (n - k_i)(n - k_{i'})/n}{n} \\ &= \frac{k_i k_{i'} + (n - k_i)(n - k_{i'})}{n^2}. \end{aligned}$$

Hence the normalized measure of hemispheric symmetry we use is

$$\bar{HS}(x_i, x_{i'}) = HS(x_i, x_{i'})/E[HS(x_i, x_{i'})]. \tag{4}$$

For each tier of each participant in each group, we computed the average of this value.

### S4. Common tier connections

Since hemispheric symmetry was not seen to contribute to hierarchical complexity, we asked instead if highly common and uncommon neighbours of Tiers explained the trends in hierarchical complexity. An ROI was defined as commonly connected to a given tier if it shared links with at least 80% of that tier's ROIs. An ROI was defined as uncommonly connected to a given tier if it shared links with at most 20%, but not none, of that tier's ROIs. The common and uncommon neighbours for each tier are shown per ROI and summed as distributions over participants in Fig S8. Common connections for Tier A and uncommon connections for Tiers B and C were also computed for 100 configuration models per participant and averaged. The results are shown in Fig S9 with Wilcoxon rank sum tests and effect sizes above each plot.

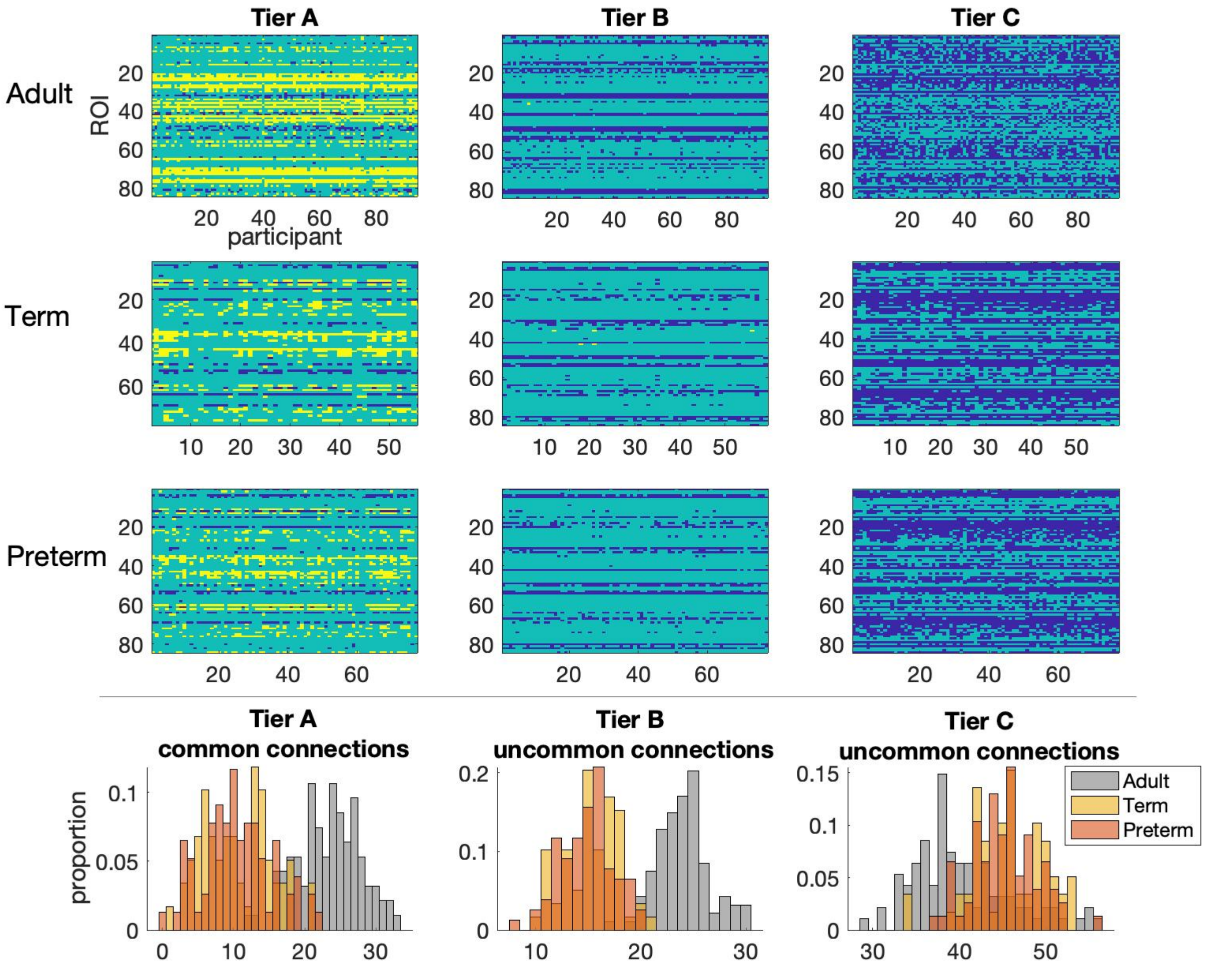


Figure S8: Top, common connections (yellow) and uncommon connections (navy blue) of the three tiers for adults, term and preterm born neonates. Axes as in the top left plot. Bottom, histograms of the number of common and uncommon connections found for each participant.

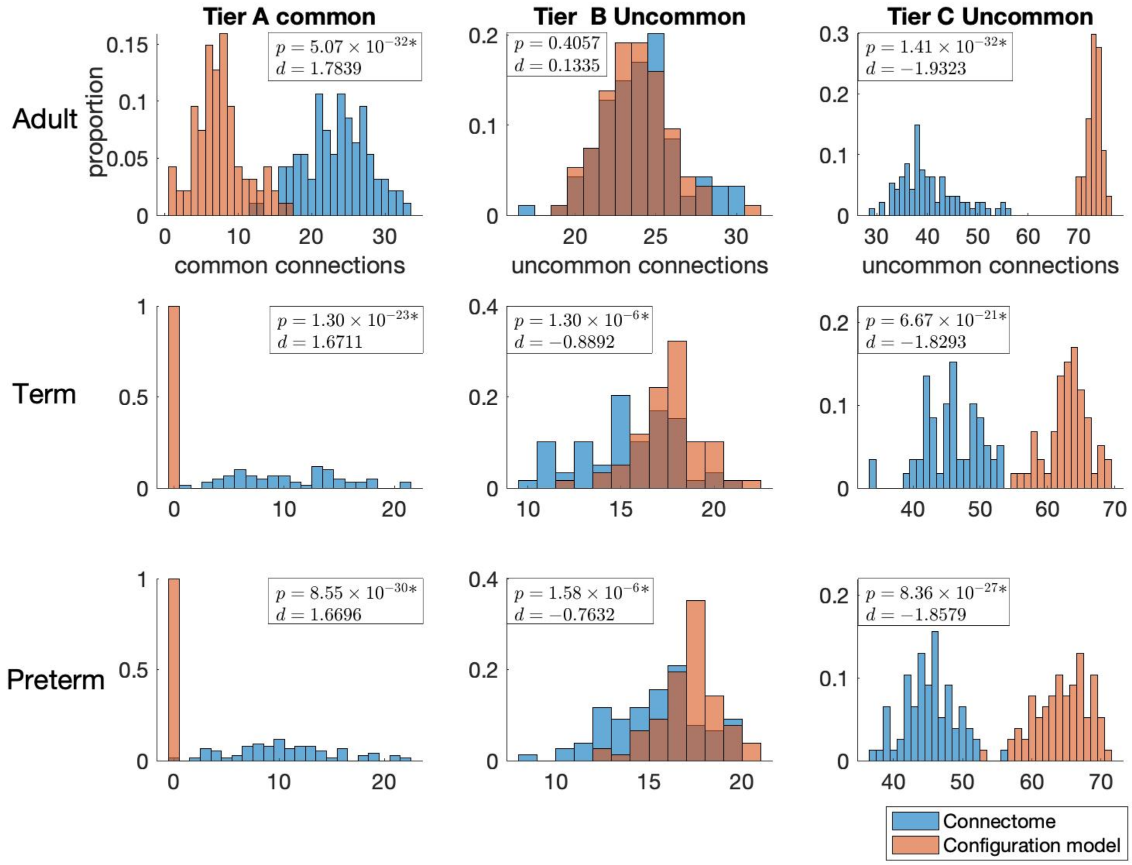


Figure S9: Tier A common connections and Tier B and C uncommon connections for adults and neonates compared against configuration models, with $p$-values from Wilcoxon rank sum tests and effect sizes (Cohen's $d$) shown. A * indicates a significant difference.

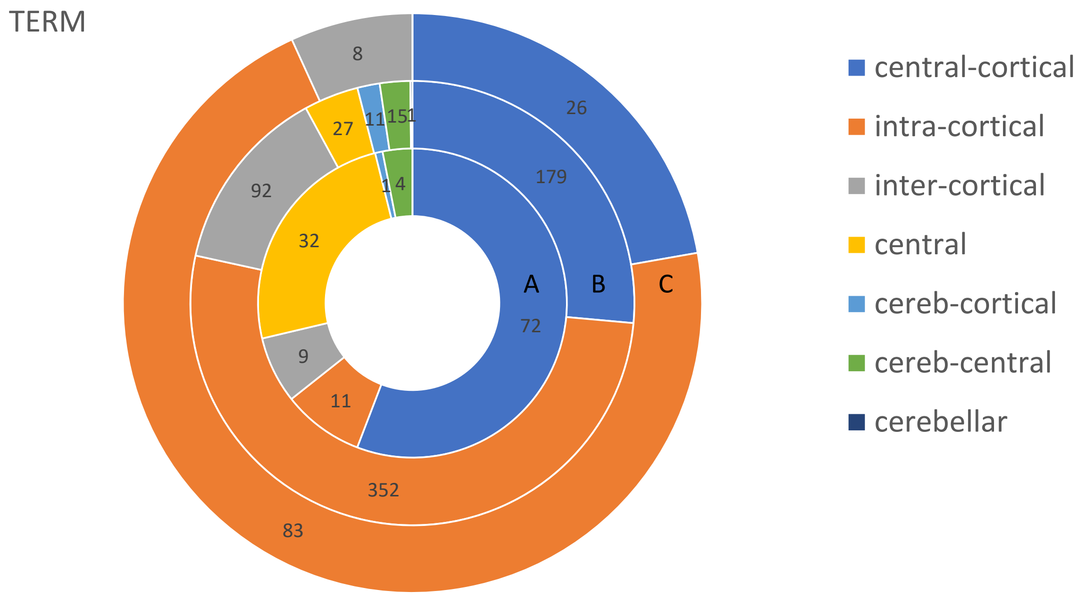


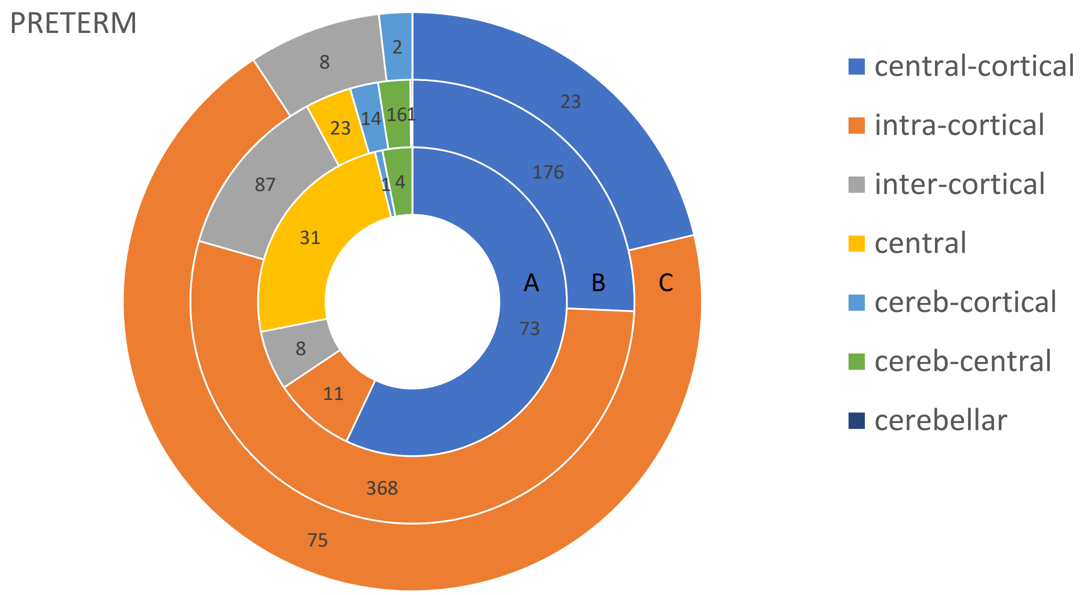


Figure S10: Distribution of connection types by tier and by group.

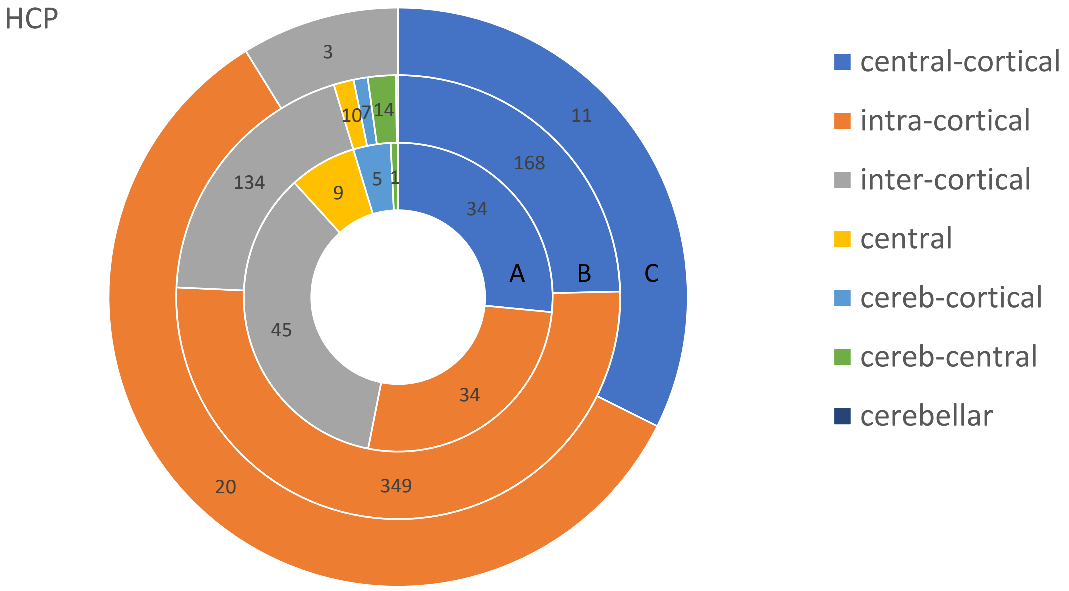


Figure S10: Distribution of connection types by tier and by group (continued).

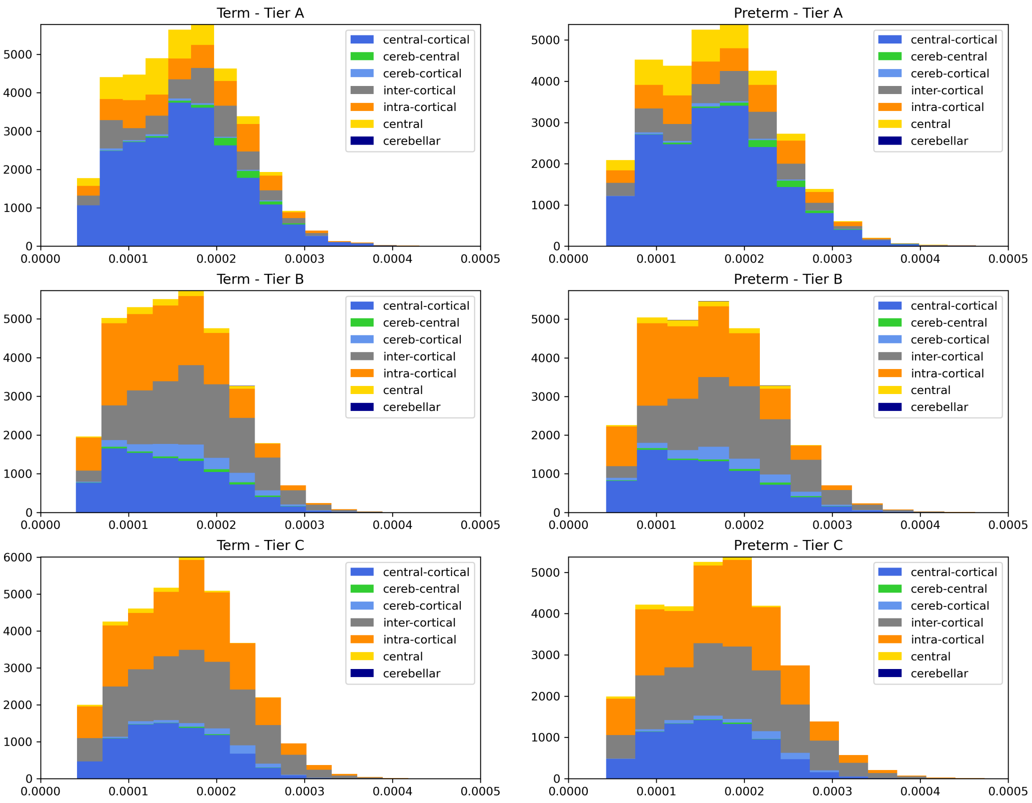


Figure S11: Histograms of connection lengths (normalized by total brain volume) by tier and by group.
